# RECLUSIVE CHANDELIERS: FUNCTIONAL ISOLATION OF DENTATE AXO-AXONIC CELLS AFTER EXPERIMENTAL STATUS EPILEPTICUS

**DOI:** 10.1101/2023.10.01.560378

**Authors:** Archana Proddutur, Susan Nguyen, Chia-Wei Yeh, Akshay Gupta, Vijayalakshmi Santhakumar

## Abstract

Axo-axonic cells (AACs) provide specialized inhibition to the axon initial segment (AIS) of excitatory neurons and can regulate network output and synchrony. Although hippocampal dentate AACs are structurally altered in epilepsy, physiological analyses of dentate AACs are lacking. We demonstrate that parvalbumin neurons in the dentate molecular layer express PTHLH, an AAC marker, and exhibit morphology characteristic of AACs. Dentate AACs show high-frequency, non-adapting firing but lack persistent firing in the absence of input and have higher rheobase than basket cells suggesting that AACs can respond reliably to network activity. Early after pilocarpine-induced status epilepticus (SE), dentate AACs receive fewer spontaneous excitatory and inhibitory synaptic inputs and have significantly lower maximum firing frequency. Paired recordings and spatially localized optogenetic stimulation revealed that SE reduced the amplitude of unitary synaptic inputs from AACs to granule cells without altering reliability, short-term plasticity, or AIS GABA reversal potential. These changes compromised AAC-dependent shunting of granule cell firing in a multicompartmental model. These early post-SE changes in AAC physiology would limit their ability to receive and respond to input, undermining a critical brake on the dentate throughput during epileptogenesis.

## Introduction

Axo-axonic cells (AACs) or Chandelier cells are an intriguing class of inhibitory neuronal subtype that uniquely target the axon-initial segment (AIS). Given the strategic location of the synapses at the primary site of action potential initiation in excitatory neurons and their expansive axon collaterals, there is considerable interest in understanding their role in shaping network activity and synchrony (Khirug *et al*., 2008; Klausberger *et al*., 2003; Varga *et al*., 2014). In the hippocampal dentate gyrus (DG), where potent inhibition to granule cells (GCs) plays an essential role in maintaining sparse firing, compromised inhibitory regulation has been proposed to contribute to the development of temporal lobe epilepsy (TLE) (Coulter and Carlson, 2007; Kobayashi and Buckmaster, 2003; Margerison and Corsellis, 1966; Zhang and Buckmaster, 2009). Consequently, several studies have examined whether AAC innervation of GCs is altered in epilepsy. Histopathological analyses of dentate tissue from human TLE patients in which AAC synapses were identified based on the characteristic synaptic cartridges have yielded variable results with reports of decrease, increase or differential patchy changes (Alhourani *et al*., 2020; Arellano *et al*., 2004; Wittner *et al*., 2001). However, dentate AACs remain an understudied population of inhibitory neurons and how their function is altered in epilepsy is not known.

A major hurdle in studying AAC function has been the paucity of unique markers to identify this neuronal subtype. While AACs share the fast spiking and non-adapting firing characteristics with parvalbumin (PV) expressing basket cells (PV-BCs), and a majority of AACs express PV, AACs are a relatively sparse population compared to PV-BCs (Buhl *et al*., 1994; Elgueta *et al*., 2015; Soriano *et al*., 1990; Taniguchi *et al*., 2013). Studies based on timed labeling of interneuron precursors have identified considerable neurochemical diversity among cortical AACs (Paul *et al*., 2017; Taniguchi *et al*., 2013). However, the strategies used to label cortical AACs do not identify hippocampal AACs, and alternative approaches show limited specificity for AACs (Ishino *et al*., 2017). Interestingly, majority of cortical AACs express the transcription factor *parathyroid hormone like hormone* (PTHLH) (Paul *et al*., 2017) suggesting it could be used to identify AACs in other circuits. AACs in distinct brain regions are known to differ in structure and physiology (Buhl *et al*., 1994; Ishino *et al*., 2017). Additionally, cortical, and hippocampal AACs show physiological differences, unique development and plasticity compared to PV-BCs (Pan-Vazquez *et al*., 2020; Rinetti-Vargas *et al*., 2017; Taniguchi *et al*., 2013). Yet, because physiological data on dentate AACs is limited and systematic comparison with PV-BCs is lacking, studies often combine fast-spiking PV interneurons during analysis. Histological data from the human epileptic DG have identified an increase in PV+ve basket-like axon terminals around GCs while PV+ve synapses at the AIS are decreased, suggesting that dentate BCs and AACs may respond differently to epileptogenesis (Alhourani *et al*., 2020). While there is considerable information on the intrinsic and synaptic physiology of PV-BC in the normal brain and in models of experimental epilepsy, whether AAC physiology is altered after status epilepticus is currently unknown (Bartos *et al*., 2001; Kobayashi and Buckmaster, 2003; Sambandan *et al*., 2010; Yu *et al*., 2013; Yu *et al*., 2016a). The diversity in physiology and seizure-induced plasticity of PV-BCs and AACs is particularly relevant due to the recent interest in activating PV neurons to curb ongoing seizure activity (Ellender *et al*., 2014; Krook-Magnuson *et al*., 2013; Ledri *et al*., 2014). Since most transgenic strategies targeting PV neurons do not distinguish between subtypes, the net effect of such interventions would depend on how both AACs and BCs inhibit GCs.

The current study was conducted to examine whether functional properties of dentate AAC are altered early after experimental status epilepticus (SE). Based on colocalization with PTHLH and the presence of axonal cartridges, we identify that a majority of PV neurons in the dentate inner molecular layer (IML) are AACs and evaluate whether dentate AACs differ from PV-BCs in their intrinsic physiological characteristics. Using a combination of single and dual patch clamp, perforated patch recordings, patterned spatial-illumination to activate optogenetically labeled PV neurons in the IML, immunostaining, and computational modeling we examine whether AAC intrinsic physiology, basal synaptic inputs, and synaptic output to GCs are altered in mice one week after pilocarpine-induced SE.

## Material and methods

### Pilocarpine status epilepticus

All procedures were performed under protocols approved by the Rutgers NJMS, Newark, NJ and the University of California Riverside, Riverside, CA, Institutional Animal Care and Use Committees. Pilocarpine injection was performed as reported previously (Zhang and Buckmaster, 2009). Young adult (6-10 weeks old), male and female C57BL/6 mice or transgenic mice with eYFP expression under PV promoter (Jackson Labs, B6;129P2-Pvalbtm1(cre)Arbr/J, 08069), were injected with scopolamine methyl nitrate (2 mg/kg, subcutaneous) 15 minutes before pilocarpine injection. Status epilepticus (SE) was induced by injection of pilocarpine (300 mg/kg, i.p). After 1h and 30 minutes of continuous stage 3 or greater seizures (Racine scale), diazepam (10 mg/kg, i.p.) was administered to terminate seizures. Age-matched control rats received scopolamine pre-treatment followed by saline injection (i.p) and diazepam after 2 hours. Additionally, few naive male and female mice were used in a subset of experiments. All anatomical and physiological studies were conducted 7-14 days after pilocarpine-SE and in age-matched, saline-injected, and naive controls.

### Slice preparation

Mice 7-14 days after saline-injection or pilocarpine-SE and age-matched naive mice were anesthetized with isoflurane and decapitated. Horizontal brain slices (300 µm) were prepared in ice-cold sucrose artificial CSF (sucrose-aCSF) containing (in mM) 85 NaCl, 75 sucrose, 24 NaHCO_3_, 25 glucose, 4 MgCl_2_, 2.5 KCl, 1.25 NaH_2_PO_4_, and 0.5 CaCl_2_ using a Leica VT1200S Vibratome (Wetzlar, Germany). The slices were incubated at 32 ± 1°C for 30 minutes in a submerged holding chamber containing 50% sucrose-aCSF and 50% recording aCSF and subsequently moved to room temperature (RT). The recording aCSF contained (in mM) 126 NaCl, 2.5 KCl, 2 CaCl_2_, 2 MgCl_2_, 1.25 NaH_2_PO_4_, 26 NaHCO_3_ and 10 D-glucose. All solutions were saturated with 95% O_2_ and 5% CO_2_ and maintained at a pH of 7.4 for 1–6 h.

### In vitro electrophysiology

For patch clamp recordings, slices were transferred to a submerged recording chamber and perfused with oxygenated aCSF at 33 ± 1°C. eYFP-positive cells were identified under epifluorescence, and whole-cell voltage- and current-clamp recordings were performed using IR-DIC visualization techniques with a Nikon AR microscope, using a 40X water-immersion objective. Recordings were obtained using Axon Instruments MultiClamp 700B (Molecular Devices, Sunnyvale, CA). Data were low pass filtered at 2 kHz, digitized using DigiData 1440A and acquired using pClamp10 at 10 kHz sampling frequency. Recordings were obtained using microelectrodes (3–5 MΩ) containing (in mM) 70 KCl, 70 K-gluconate, 10 HEPES, 2 MgCl_2_,

0.2 EGTA, 2 Na-ATP, 0.5 Na-GTP and 10 PO Creatine titrated to a pH 7.25 with KOH in the absence of synaptic blockers. Biocytin (0.2%) was included in the internal solution for post-hoc cell identification (Yu *et al*., 2013; Yu *et al*., 2016a). Recorded neurons were initially held at -70 mV and the response to 1.5 sec positive and negative current injections were examined to determine active and passive characteristics. Post-hoc biocytin immunostaining and morphological analysis were used to definitively identify AACs included in this study, based on the presence of axonal cartridges for all cells in Fig 1. For data presented in Fig 2-4, AACs were targeted as PV/YFP labeled neurons in the IML and confirmed based on post hoc morphological recovery of widespread dendritic arbors and presence of axonal cartridges in the granule cell layer (Supplementary Fig. 2). Following current-clamp recordings, cells were held in voltage-clamp at -70 mV for analysis of GABA currents and sEPSCs were blocked by kynurenic acid 3mM. In experiments where sEPSCs were recorded, saturating concentrations of the GABA_A_R antagonist SR95531 (10 µM) were included.

**Figure 1:**
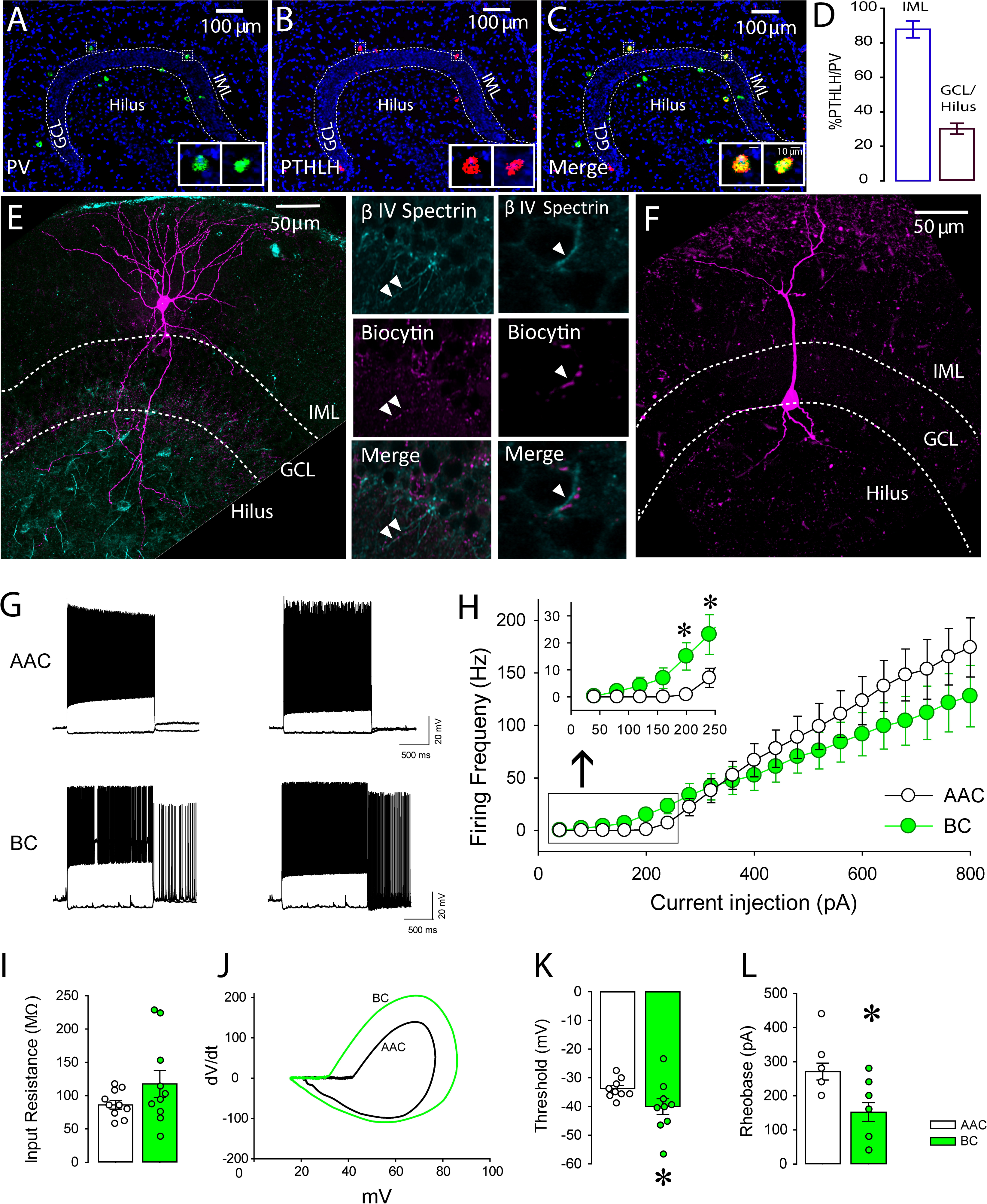
Characterization of dentate parvalbumin-positive BCs and AACs. (A-C) Epifluorescence images of the dentate gyrus illustrate localization of probes for PV in green (A) PTHLH in red (B) and merged image showing co-labeling (C). Insets illustrate magnified images of PTHLH and PV co-labeled neurons in the inner molecular layer (IML). (D) Summary graph of percentage of PTHLH/PV positive neurons in IML vs. Hilus. (E-F) Maximum intensity projections of confocal image stacks of a representative AAC (E) BC (F). Images to the right of (E) illustrate two examples of axons from the cell in E showing close apposition for βIV-spectrin labeling for axon initial segments (top), and biocytin (middle) as shown in merge (bottom). Arrowheads indicate regions of close apposition between βIV-spectrin labeling on axon initial segment of GC and biocytin filled axonal cartridge of AAC. (G) Representative suprathreshold traces from two different AAC (top) and BC (bottom) neurons showing persistent firing in BCs but not in AACs. (H) The average firing frequency of BCs and AACs in response to increasing current injection shows differences in firing frequency at low amplitude current injections (inset). (I) Summary plots of input resistance in BC and AAC. (J-L). Example phase plots (J) and summary of action potential threshold (K) and, rheobase (LK) in BCs and AACs. Scale: 100μm. * indicates p<0.05 by unpaired Student’s t-test. Note: GCL – Granule Cell Layer, IML – Inner Molecular Layer.

**Figure 2:**
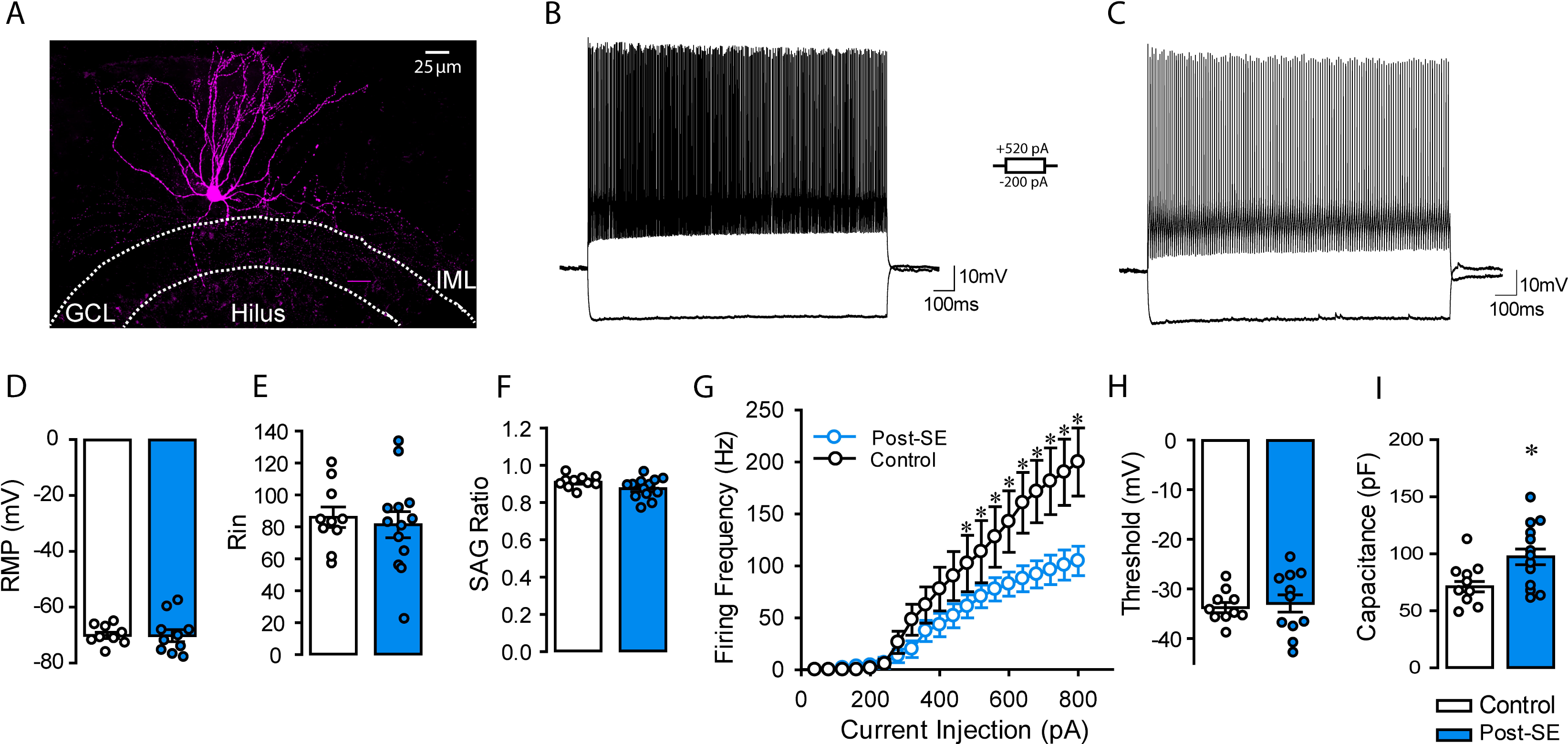
**AAC excitability is compromised after status epilepticus**. (A) Representative confocal image of AAC in the dentate IML region and spanning axonal cartridges. (B-C) Membrane voltage traces of axo-axonic cells from control (B) and post-SE mice (C) during a 520 pA and -120 pA current steps. (D-F) Summary histograms of resting membrane potential (D), input resistance (E), and SAG ratio (F) from control and post-SE groups. (G) Overlay of the firing rates of control and post-SE AACs in response to increasing current injections. (H) Summary graph of threshold in AACs from control and post-SE mice. (I) Summary plot of AAC membrane capacitance * in panel I indicates p<0.05 by unpaired Student’s t-test and in panel G represents p<0.05 by two-way repeated measure ANOVA.

### Paired recordings and optically evoked responses

Unitary inhibitory postsynaptic (uIPSC) responses were evoked in the dentate GCs (voltage clamped at -70mV) by optically stimulating presynaptic ChR2 expressing cells using a Digital Mirror Device (DMD)-based pattern illuminator (Mightex Polygon 400), coupled to 473nm blue LED (AURA light source), and controlled via TTL based input from pClamp. A circular ROI of ∼ 50μm diameter was marked around 1 to 2 eYFP expressing putative AACs in the IML. In a subset of experiments, optical activation (5 ms) of putative AACs recorded in the IML reliably elicited single action potentials (not shown). Light power was fixed to elicit single uIPSCs in the recorded postsynaptic GC and recordings without postsynaptic responses were excluded from analysis. Alternatively, eYFP-expressing neurons in the IML were targeted using microelectrodes containing KCl-KGluc based internal solution. Selected ROIs were stimulated by a brief 5ms single light pulse or train of 3 ms light pulses at 20Hz and repeated for 10 recording sweeps with an inter sweep interval of 10s.

### Gramicidin perforated patch recording

Granule cells were recorded using Gramicidin-perforated patch to measure the equilibrium potential of GABA (*E_GABA_*) with intact intracellular chloride concentration. Patch electrodes were filled with KCl-K-gluconate-based internal solution with gramicidin (20 μg/mL, G5002, Merck KGaA), 5-CFDA, AM (40 μM, C1354, Invitrogen, Rinetti-Vargas *et al*., 2017), and Alexa Fluor™ 594 Hydrazide (1 μM, A10438, Invitrogen, Hsu *et al*., 2019). After 20-30 minutes in the cell-attach configuration, perforated patch was confirmed by an access resistance of 70∼100 MΩ and the presence of 5-CFDA, AM but absence of Alexa Fluor™ 594 Hydrazide in the recorded soma. The AIS-mediated GABA currents were evoked by focal pressure ejection of GABA_A_R agonist muscimol (100 μM, 0289, TOCRIS) toward the AIS of recorded GCs via a glass pipette controlled by a Pneumatic PicoPump (100 ms, PV 800, World Precision Instruments). The relation of evoked current amplitude and holding potential was plotted to determine *E_GABA_* for each cell.

### Anatomical methods

To visualize PTHLH and PV spatial colocalization, we used RNAscope *Multiplex Fluorescent In situ hybridization* (*m-FISH*) on cryosectioned (50 µm) tissue from control and mice one-week post-SE. RNAscope Multiplex Fluorescent Reagent Kit v2 Assay was used following the protocol for fixed frozen tissue (Nguyen *et al*., 2020; Wang *et al*., 2012). The following kits and probes were used for this study: RNAscope® Multiplex Fluorescent Reagent Kit (Cat. #323100), RNAscope® Probe-Mm-Pthlh (Cat. #456521), and RNAscope® Probe-Mm-Pvalb-C2 (Cat No. 421931-C2). Slides were coverslipped using Vectashield mounting media (Vector Labs) and imaged using a Zeiss Axioscope and analyzed using Stereo Investigator (Microbrightfield).

Following physiological recordings, slices were fixed in 0.1 M phosphate buffer containing 4% paraformaldehyde at 4°C for two days or fixed briefly for 30 minutes for staining with anti-βIV-spectrin. For post-hoc immunohistochemistry, thick slices (300 µm) were incubated overnight at room temperature with either of these following primary antibodies: anti-parvalbumin antibody (PV-28, 1.5:1000, polyclonal rabbit, Swant), anti-GFP antibody (AB 16901, 1:500, polyclonal chicken, Millipore) and anti-βIV-spectrin (AB2315816, 1:500, monoclonal mouse, NeuroMab) in 0.3% Triton X-100 and 3% normal goat serum containing PBS. Immunoreactions were revealed using Alexa 488-conjugated secondary goat antibodies against rabbit IgG (1:250), and biocytin staining was revealed using Alexa 594-conjugated streptavidin (1:1000). Sections were visualized and imaged using a Nikon A1R laser confocal microscope with a 1.2 NA 60X water objective.

Immunohistological studies were performed in mice one week after SE and in age-matched controls. Briefly, mice were anesthetized with isoflurane and sacrificed by decapitation, brains were rapidly removed and fixed by immersion fixation in 4% paraformaldehyde overnight at 4°C. Sections were cut at 50 µm thickness on a Vibratome, and every sixth section was selected from the entire septotemporal extent of the hippocampus. Sections were processed for double immunofluorescence labeling for PV and eYFP. NeuN immunofluorescence was used to assess cell loss. Briefly, free-floating sections were rinsed in PBS, were blocked with 10% NGS and 0.3% Triton X-100 in PBS at RT for 1 h and then incubated in a solution containing monoclonal mouse anti-PV (1.5:1,000; 235, Swant), polyclonal chicken anti-GFP (AB 16901, 1:500, Millipore) or mouse anti-NeuN (MAB377, 1:10,000, Millipore) in PBS with 0.3% Triton X-100 and 3% NGS at RT for 24 h. Sections were then rinsed in PBS and incubated at 4°C for 24 h in a mixture of goat anti-mouse IgG conjugated to Alexa Fluor 594 (1:500) and goat anti-chicken IgG conjugated to Alexa Fluor 594 (1:500) to reveal PV and eYFP respectively. Similarly, goat anti-mouse IgG conjugated to Alexa Fluor 488 (1:500) was used to reveal NeuN. Sections were rinsed the next day in PBS and mounted with Vectashield (Vector Labs). Controls in which primary antibody was omitted were routinely included.

For a subset of experiments mice one week after SE and age-matched controls were transcardially perfused with cold PBS followed by cold 4% paraformaldehyde (PFA). Brains were incubated in 4% PFA overnight, switched to 30% sucrose for 2-3 days to allow tissue to sink, and were flash frozen in OCT using liquid nitrogen and stored at -80C. Tissue was cryo-sectioned at 20 µm using a Leica Cryostat, mounted on SuperFrost slides, and stored at -80C prior to staining. Co-staining for perineuronal nets and PV was performed as described previously (Brewton *et al*., 2016). Tissue was washed with PBS, quenched with 50 mM NH_4_Cl for 15 min at RT, permeabilized with 0.1 % Triton X-100 for 10 min and blocked with 5 % NGS and 1 % BSA. After blocking, tissue was incubated in primary antibody solution containing monoclonal mouse anti-PV (1:1000, #235, Swant) and FITC-conjugated Wisteria Floribunda Agglutinin (WFA, 1:500, FL-135, Vector Laboratories) overnight at 4 ℃. Tissue was then washed and incubated in secondary antibodies (1:500, goat anti-mouse IgG conjugated to Alexa Fluor 647) for 1hr at RT, washed, and coverslipped with Vectashield with DAPI (Vector Labs). Slides were imaged using Zeiss 880 Upright Airyscan Microscope at 63x and images were analyzed using ImageJ by drawing an ROI with WFA staining around PV neurons in the IML and excluding the unstained region in the center. Since WFA staining was variable across sections, the integrity of perineuronal nets was quantified as the number of peaks in intensity over the average intensity in the ROI (Tewari *et al*., 2018). Data are presented normalized to the area of the ROI. For examination of AIS length and GABA_A_ receptor α2 subunit staining, tissue was blocked with 10% NGS and 0.3% Triton X-100 at RT for 1h then incubated in primary antibodies for AnkG (1:500, Cat#386-006, Synaptic Systems, Gao and Heldt, 2016) and GABA_A_ α2 (1:1000, Cat#224-103, Synaptic Systems) in 5% NGS PBST overnight at 4°C. Tissue was then washed and incubated in secondary antibodies for 1hr at RT, washed, and coverslipped with Vectashield with DAPI (Vector Labs). Slides were imaged using Zeiss 880 Upright Airyscan super resolution at 63x and AIS length and puncta count was performed using QuPath 0.4.2 software (https://qupath.github.io). Neurolucida 360 was used to reconstruct AIS and GABA_A_ α2 puncta for representative images.

Cell counts were performed using the Optical Fractionator probe of Stereo Investigator V.10.02 (MBF Bioscience) using an Olympus BX51 microscope and a 40X objective by an investigator blinded to treatment as previously described (Gupta *et al*., 2012). In each section, the hilus was outlined by a contour traced using a 10X objective. The following sampling parameters were set at 40X: counting frame, 100 µm X 100 µm; dissector height, 45 µm; and guard zone distance set at 5µm. Approximately 25 sites per contour, selected using randomized systematic sampling protocols, were sampled. In each section, the cell count was estimated based on planimetric volume calculations in Stereo Investigator.

### Nonstationary fluctuation analysis

Nonstationary variance analysis (Sigworth, 1980; Yu *et al*., 2016b) was performed on unitary postsynaptic current responses from GCs that were optically evoked using DMD-based stimulation at single cell resolution. Stable recordings were pooled from individual groups to isolate fluctuations in the current decay attributable to stochastic channel gating, the mean waveform was scaled to the peak of individual IPSCs (De Koninck and Mody, 1994; Traynelis *et al*., 1993). Successful responses were used to calculate the ensemble mean current (I) and peak-scaled variance (σPS2) for each data point. Plots of variance versus current were fit with the equation: σPS^2^ = iI − (I^2^/NP) + σB^2^ , where i is the weighted-mean single-channel current, NP is the number of channels open at peak synaptic current, and σB 2 is the background variance (Brickley *et al*., 1999). Only IPSCs that showed stable peak amplitude over time were included in the analysis.

### Computational Model

A multicompartmental granule cell model based on Santhakumar *et al*. (2005) was simulated using NEURON 7.8.2 (Hines and Carnevale, 1997) on a PC run with windows 10. A single granule cell (GC) multicompartmental model (Santhakumar *et al*., 2005; Yu *et al*., 2013) was modified to include an axon initial segment (AIS) of length (30 μm) and diameter (2 μm) connected to the somatic compartment. The AIS compartment was modeled to include sodium, fast and slow delayed rectifier potassium and leak channels which were correspondingly decreased in the somatic compartment (See Supplemental Table 1). Model GC included two AMPA and two GABA synapses on the middle dendritic and AIS segments, respectively, implemented using Exp2Syn mechanism. The rise, weighted tau decay, and reversal potential for excitatory AMPA synapses are 1.5 ms, 5.5 ms, and 0 mV respectively, while inhibitory GABA synapses were modelled with rise, decay, and reversal potential of 0.2 ms, 6-10 ms and, -65 mV (constrained by experimental data). A single presynaptic stimulus from an artificial cell is used to drive two AMPA synapses located in the middle dendritic compartments of the model GC. Increasing peak excitatory conductance in the range of 10-30 nS at each AMPA synapse were simulated. Based on experimental estimates of AIS GABA synapse peak conductance obtained from non-stationary fluctuation analysis, increasing peak GABAergic conductances from 0.5 nS – 10 nS in each of the two AIS GABA synapses were evaluated. Results are reported as ‘1’ representing action potential and ‘0’ as shunted action potential.

### Analysis and statistics

Analysis was performed by a researcher blinded to treatment groups, followed by unblinding to categorize treatment groups for statistical analysis. Synaptic currents were measured as described previously (Gupta *et al*., 2012; Yu *et al*., 2013; Yu *et al*., 2016a) using custom macros in IgorPro7.0 software (WaveMetrics, Lake Oswego, OR). Recordings were discontinued if series resistance increased by >20%. Individual synaptic events were detected using the custom software in IgorPro7.0 (Gupta *et al*., 2012; Santhakumar *et al*., 2010). Events were visualized, and any “noise” that spuriously met trigger specifications was rejected. Cumulative probability plots of sIPSC and sEPSC parameters were constructed using IgorPro by pooling an equal number of sIPSCs and sEPSCs from each cell. Passive parameters such as input resistance and SAG ratio were analyzed from hyperpolarizing current steps (-200pA). Membrane time constant was estimated from a standard single exponential fit to voltage responses to a -200pA hyperpolarizing step using the Levenberg-Marquardt method in Clampfit. Cell capacitance was calculated as the ratio of membrane time constant and input resistance of cell. Active properties were estimated from the depolarizing current steps. The threshold for the action potential was determined by calculating the first time derivative (*dV/dt*) of the voltage trace and setting 30mV/ms as the threshold for level for action potential initiation (Gupta *et al*., 2012). Rheobase was determined as the minimum current input needed for the cell to cross the threshold and fire an action potential. Data that were over two standard deviations from mean were excluded. Decay kinetics from uIPSC traces were calculated based on single exponential fit on the decay phase of mean uIPSC trace from peak to baseline. Statistical analysis was performed by paired and unpaired Student’s *t*-test (Microsoft Excel 2007), or Mann Whitney U-test for data that were not distributed normally or univariate and multivariate repeated measures ANOVA (Sigma Plot 12.3) for experiments involving repeated measurements from the same sample. Significance was set to *p* < 0.05. Data are shown as mean ± s.e.m or median and inter-quartile range (IQR) where appropriate.

## Results

### Preferential localization to the inner molecular layer and higher rheobase distinguishes dentate axo-axonic cells from basket cells

Since the DG circuit function is heavily regulated by inhibition, and AACs are ideally poised to control GC output, there has been considerable interest in evaluating dentate AAC plasticity in epileptic networks. Recent analyses of cortical AACs have revealed distinctive laminar distribution and connectivity, and identified transcriptional markers enriched in AACs (Paul *et al*., 2017). We utilized the expression of *parathyroid hormone like hormone* (PTHLH), a transcription factor shown to be selectively expressed in over 95% of cortical AACs (Paul *et al*., 2017), to identify the distribution of dentate AACs. Using *RNAscope in-situ hybridization* for PV and PTHLH, we identified PV and PTHLH colabeled somata in the dentate IML, granule cell layer (GCL) and hilus. Layer specific analysis revealed significantly fewer PV+ve interneurons (PV-INs) in the IML compared to the GCL/hilus (Figure 1A-C, cells/section, GCL/hilus: 7.5±0.6; IML: 2.58±0.16, 5-9 sections each/4 mice, p=0.0002 by *Student’s t*-test). PTHLH expressing neurons were present in the IML and the GCL/hilus with over 75% of the PTHLH+ve neurons colabeling for PV. Notably, majority of the PV neurons in the IML colocalized PTHLH indicating that they are likely to be AACs, while fewer GCL/hilar PV-INs co-localized PTHLH (Figure 1D, % of PV-INs positive for PTHLH, IML: 87.9±4.9, GCL/hilus: 29.8±3.2, 5-9 sections each/4 mice, p<0.0001 by *Student’s t*-test). These data suggest that PV+ve neurons in the DG IML are likely to be AACs.

To determine whether IML PV neurons show the morphological features of AACs and evaluate whether dentate AACs are physiologically distinct from PV-BCs in the GCL/hilar border, we undertook whole cell recordings from labeled neurons in PV-reporter mice. Specificity of the PV-ChR2::YFP reporter was confirmed by expression of eYFP in ∼80% of PV+ve neurons and colocalization of PV in all eYFP positive cells (Supplementary Fig. 1). PV-cre mice that were not crossed with reporter lines and cre-negative mice lacked eYFP expression (data not shown), validating the specificity of transgenic mice. eYFP positive neurons with somata in IML and GCL-hilar border were recorded under IR-DIC visualization, filled with biocytin, and processed for post hoc immunostaining to recover axonal arbors and distinguish AACs from basket cells. Consistent with the expression of PTHLH in AACs, PV+ve IML neurons in which axons were recovered consistently showed the presence of vertical axonal cartridges (n=10 cells, Supplementary Fig. 2) with 5/5 cells tested showing close apposition of AAC axonal cartridges with the AIS labeled with βIV-spectrin (Figure 1E right panels). In contrast, axons of PV-INs in the GCL/hilar border showed perisomatic projections typical of basket cells and did not contact the AIS (Figure 1F). Morphologically identified AACs had multipolar somata in the IML and, typically, extended prominent apical dendrites reaching fissure and a few had basal dendrites in the hilus suggesting that AACs could contribute to both feedforward and feedback GC inhibition. 3D confocal imaging revealed extensive dendritic arborization along the z-plane distinguishing the dendritic structure of AACs from semilunar granule cells in the IML (Afrasiabi *et al*., 2022; Gupta *et al*., 2020).

While prior studies have demonstrated that dentate AACs show the fast-spiking characteristic of PV-INs (Buhl *et al*., 1994; Elgueta *et al*., 2015), systematic comparison of dentate AACs and BCs intrinsic physiology is currently lacking. Both morphologically identified AACs in the IML, and BCs in the GCL/hilar border displayed the characteristic fast-spiking pattern with little adaptation (Figure 1G). AAC firing frequency in response to 200-240 pA depolarizing current injection (I_inj_) was significantly lower than that of BCs (Figure 1H, AP frequency at I_inj_ 200pA in Hz: BCs: 15.0±5.1, AACs: 0.8±0.6 and I_inj_ 240pA, BCs: 23.1±7.3, AACs: 7.0±3.5; n=9 AACs and n=10 BCs, unpaired t-test: p<0.05). However, AACs trended to fire at higher frequency than BCs with increasing I_inj_ and rarely showed depolarization block which was observed in BCs. Consequently, the overall firing frequency in response to increasing current injections was not different between cell types (two-way RM ANOVA). Dentate PV-INs have been shown to exhibit persistent firing in the absence of current input following depolarization (Elgueta *et al*., 2015). Interestingly, while BCs (5 / 10 cells) developed persistent baseline firing following the depolarizing current injections, none of the AACs (n=10) exhibited persistent firing (Figure 1G). Comparison of passive membrane properties revealed no difference in resting membrane potential (RMP in mV, AAC: -70.12±3.42; BCs: -68.58±2.42, n=9 each, unpaired t-test p>0.05), input resistance (Figure 1I, R_in_ in MΩ, AACs: 86.1±6.5; BCs: 117.4±20.3, n = 10 each, unpaired t-test p>0.05) and SAG ratio (AACs: 0.91±0.01; BCs: 0.85±0.03, n = 10 each, unpaired t-test p>0.05) between AACs and BCs. However, the threshold for action potential firing in AACs was significantly more depolarized than in BCs (AP threshold in mV, AACs: -33.7±1.0; BCs: - 40.0±2.7, n = 10 each, unpaired t-test p<0.05) (Figure 1J, K). Consistently, rheobase, a measure of minimum current needed to evoke action potential, was higher in AACs compared to BCs (AACs: 271.1±24.7 pA; BCs: 152.0±27.8 pA, n = 9 & 10 respectively, unpaired t-test p<0.05) (Figure 1L). These data demonstrate that PV+ve neurons in the dentate IML are predominantly AACs and share the high-frequency non-adapting firing characteristics with BCs, but, are distinguished by higher threshold to activation and greater reliability/fidelity during sustained activity.

### AACs survive status epilepticus and are less excitable

Studies in human and experimental epilepsy have identified changes in the PV+ve axonal cartridges in the DG, raising the possibility that AACs are lost or structurally altered during epileptogenesis (Alhourani *et al*., 2020; Arellano *et al*., 2004; Ribak, 1985; Wittner *et al*., 2001). We implemented the pilocarpine-induced status epilepticus (Kobayashi and Buckmaster, 2003; Peng *et al*., 2004; Zhang and Buckmaster, 2009), to examine early changes in AACs during epileptogenesis. NeuN labeling for neuronal somata confirmed a significant decrease in hilar neurons one week after SE (Supplementary Fig. 3, neurons/section, Control: 154.6±9.2 from 17 sections/3 mice; Post-SE: 109.3±10.8 from 15 sections/ 3 mice, *p<* 0.05, unpaired t-test), consistent with earlier studies (Kobayashi and Buckmaster, 2003; Mello *et al*., 1992; Yu *et al*., 2013). While there was a trend towards a decrease in PV expressing neurons in the hilus one week Post-SE (Supplementary Fig. 3, neurons/section, Control: 16.9±2.2 in 11 sections/3 mice; Post-SE: 12.8±1.6 in 17 sections/3 mice, *p>*0.05, unpaired t-test), PV-IN counts in the IML were not altered after SE (Supplementary Fig. 3, neurons/section, Control: 6.7±1.3 in 11 sections/ 3 mice; Post-SE: 10.2±1.4 in 17 sections/3 mice, *p>* 0.05, unpaired t-test) indicating that the AACs in the IML survive SE.

We examined the intrinsic physiology of AACs in the IML from mice one to two weeks after status epilepticus (post-SE) and age-matched saline injected controls in the absence of synaptic blockers. AAC passive properties including resting membrane potential (RMP in mV, Control: - 70.12±3.42, n=9; Post-SE: -70.22±6.85, n=12, unpaired t-test p>0.05), input resistance (R_in_ in MΩ, Control: 86.1±6.4, n= 10; post-SE: 81.3±8.2, n=13, unpaired t-test p>0.05), membrane time constant (in ms, Control: 6.84±0.52, n=10; post-SE: 8.18±0.63, n=12, unpaired t-test p>0.05) and sag ratio (Control: 0.91±0.01, n=10; post-SE: 0.87±0.01, n=12, unpaired t-test p>0.05) were not altered after SE (Figure 2A-F). Additionally, slow AHP, calculated as the difference between baseline and the lowest anti-peak amplitude at the end of spike train in response to a 520pA current step was not different between groups (Control: -1.66±0.67, n=10; post-SE: -3.89±0.54, n=12, unpaired t-test, p=0.4). However, AACs in post-SE mice showed a consistent reduction in firing rate during somatic current injection with a significant effect of treatment (Figure 2G, F_(1,19)_= 5.16, p<0.05 by Two-way RM ANOVA) and significant interaction between treatment and I_inj_ (F_(1,19)_= 6.73, p<0.05 by Two-way RM ANOVA). AAC firing in response to I_inj_ >500 pA was consistently reduced in post-SE mice (Figure 2G, Two-way RM ANOVA with Bonferroni corrected pairwise comparison). However, action potential threshold (Figure 2H, Control: - 33.7±1.0 mV, n= 10; post-SE: -32.9±1.7 mV, n=12, unpaired t-test p>0.05) and rheobase (Control: 271.1±24.7 pA, n=10; post-SE: 333.3±47.05 pA, n=12, unpaired t-test p>0.05) were not different between groups. Because post-SE AAC firing was reduced despite a lack of change in threshold or rheobase, and previous studies had identified that post-seizure changes in neuronal capacitance could impact excitability (Tewari *et al*., 2018; Whitebirch *et al*., 2022) we compared the membrane capacitance (C_m_) between AACs in control and post-SE groups. AAC capacitance was significantly increased after SE (Figure 2I, C_m_ in nF, Control: 71.2±4.7, n=14; post-SE: 97.3±6.9, n=16, unpaired t-test p<0.05). Since seizure induced changes in extracellular matrix have been reported to alter C_m_ (Tewari *et al*., 2018), we examined integrity of the perineuronal net around PV neurons in the dentate IML by staining for Wisteria Floribunda Agglutinin (WFA) which labels the perineuronal net. Consistent with an increase in AAC capacitance, there was a significant decrease in the number of WFA intensity peaks around IML PV neurons after SE (Supplementary Fig. 4). Overall, these data identify a post-SE reduction in the ability of AACs to sustain high firing frequency which contrasts with the lack of change in PV-BC intrinsic physiology after SE (Yu *et al*., 2013).

### SE reduces synaptic inputs to AACs

Since SE leads to early changes in dentate neuronal networks (Kobayashi and Buckmaster, 2003; Peng *et al*., 2013; Yu *et al*., 2016a), we examined whether baseline synaptic inputs to morphologically identified AACs were altered after SE. Spontaneous excitatory postsynaptic currents (sEPSCs) in AACs were recorded from a holding potential of -60 mV in the presence the GABA_A_Rs antagonist gabazine (10µM). Compared to age-matched controls, the frequency of sEPSCs in AACs was reduced after SE, although the effect size was modest (Figure 3A, B; Control: Median=12.1Hz; IQR=5.6-21.3, n=8; Post-SE: Median=6.8Hz; IQR=2.5-23.8, n=6 by Mann-Whitney U-test, Cohen’s D: 0.37). However, there was a robust increase in sEPSC amplitude in AACs from post-SE mice, with a moderate effect size, (Figure 3A, C; Control: Median=19.4 pA; IQR=15.1-25.1, n=8; Post-SE: Median=29.1 pA; IQR=21.1-39.2, n=6 by Mann-Whitney U-test; Cohen’s D: 0.89). The rise time and τ_Decay_of AACs sEPSCs were not altered after SE (Supplementary Fig. 5A-B).

**Figure 3:**
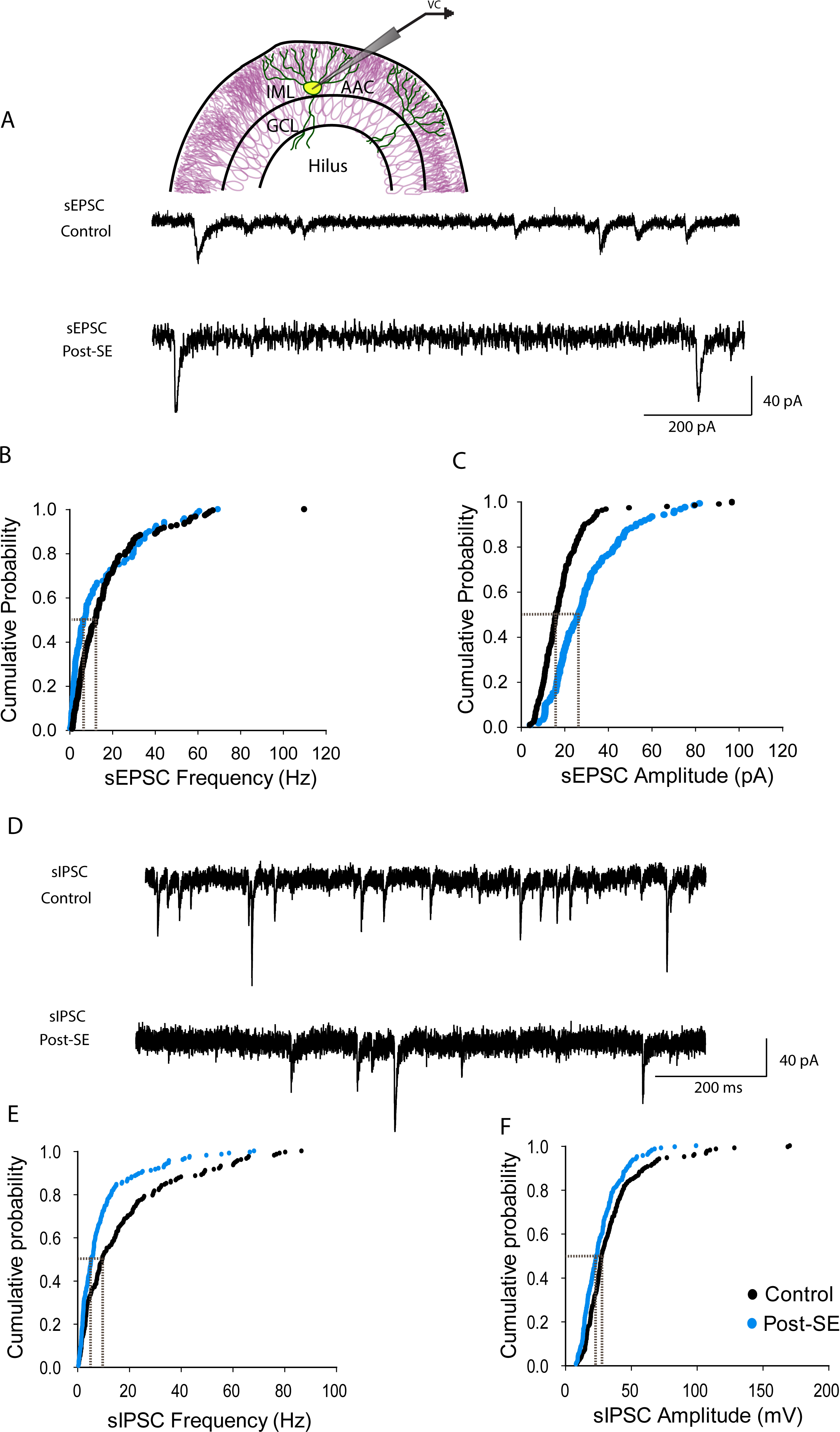
Status epilepticus alters synaptic drive to dentate AACs in the inner molecular layer. (A) Schematic of recording configuration illustrates targeting eYFP positive IML-AAC neurons (above) Representative current traces show spontaneous excitatory postsynaptic currents from AACs in control (top) and post-SE mice (bottom). (B-C) Cumulative probability plots of sEPSC frequency (B) and amplitude (C) recorded in gabazine (10µM) (inset gray lines shows differences at 50% probability). (D) Example traces show spontaneous inhibitory postsynaptic currents in AACs from control (top) and post-SE (bottom) mice. (E-F) Cumulative probability plots of sIPSC frequency (E), and amplitude (F), recorded in kynurenic acid (3mM) (inset gray dotted lines shows differences at 50% probability). * Indicates p<0.05, by Mann-Whitney U Test, n=10 cells each.

**Figure 4:**
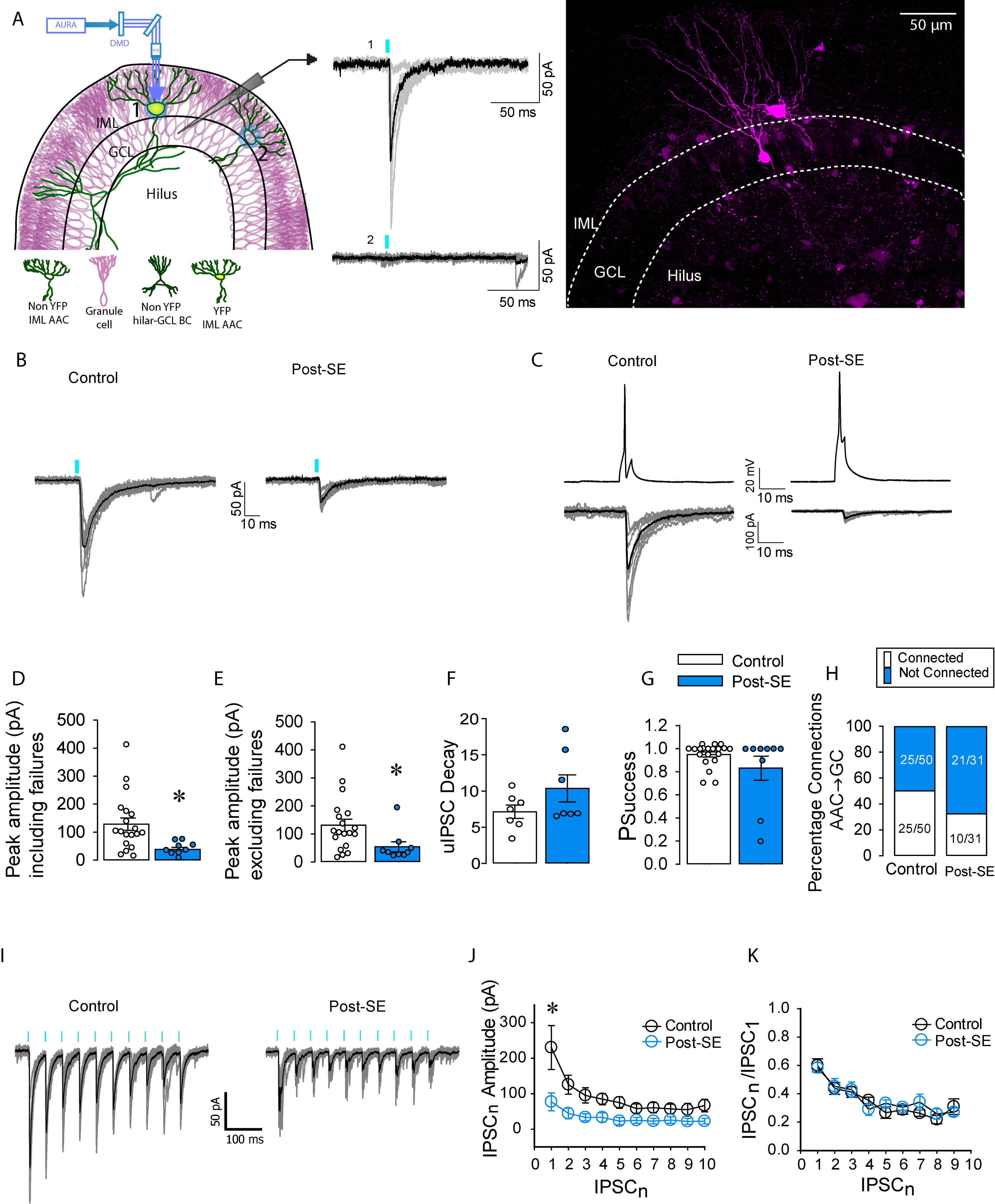
Decreased synaptic strength at AAC-GC synapses early after SE. (A) Schematic of dentate gyrus slice on the left illustrating voltage-clamped granule cells, and typical eYFP+ AAC and eYFP-neuron in the IML region indicated by 1 and 2, respectively. Line diagram in blue shows light path routed through the DMD. Representative traces on the right indicated by 1 show control experiments where DMD stimulation of eYFP+ neuron evokes uIPSCs in coupled GC, whereas DMD stimulation of eYFP-neuron failed to elicit responses in GC indicated by 2. Example confocal image on the right shows biocytin-filled GC and AAC filled during paired recording. (B) Representative uIPSCs evoked (10 sweeps in gray) in synaptically coupled GCs by discharging single spikes in presynaptic AACs from control (left) and post-SE (right) mice. Average uIPSC traces are in black. Blue bar indicates light activation. (C) Postsynaptic responses in coupled GC in control (left) and post-SE (right) elicited by presynaptic action potentials (above) in AAC. (D-G) Summary histograms for AAC mediated uIPSC peak amplitude including failures (D), excluding failures (E), weighted decay (F) and probability of success (G). (H) Summary of percentage of synaptic connections from AAC to GC. (I) Representative traces illustrate multi-pulse depression of AAC evoked uIPSCs in GCs during optogenetic stimulation (blue bar) in control (left) and post-SE (right). The 10 individual sweeps are in gray, and the average trace is illustrated in black. (J-K) Summary plot of peak amplitude of uIPSCs elicited during the stimulus train (J) and amplitude ratio between uIPSCn and uIPSC_1_ (K). * Indicates p<0.05 by Two-way ANOVA. Note: GCL – Granule Cell Layer, IML – Inner Molecular Layer.

Next, we evaluated spontaneous inhibitory inputs to AACs held at -70 mV in the presence of glutamate receptor antagonist kynurenic acid (3 mM). The frequency of spontaneous inhibitory postsynaptic currents (sIPSCs) in AACs was significantly lower after SE (Figure 3D, E; Control: Median=14.5Hz; IQR=6.8-30.2, n=7; Post-SE: Median=6.1Hz; IQR=2.4-11.4, n=8, p<0.05 by Mann-Whitney U-test, Cohen’s D: 0.75). Similarly, there was a post-SE reduction in sIPSC amplitude after SE (Figure 3D, F; Control: Median=27.2 pA; IQR=21.0-37.7, n=7; Post-SE: Median=23.3 pA; IQR=15.4-35.0, n=8, p<0.05 by Mann-Whitney U-test, Cohen’s D: 0.35). sIPSC rise time and τ_Decay_were not different between AACs from control and post-SE mice (Supplementary Fig. 5E-F).

In a subset of paired recordings from AACs, we identified reciprocal electrical coupling between AACs (2 control and 1 post-SE AAC pair, Supplementary Fig. 6), although none of the recorded pairs showed synaptic coupling. Given the sparse distribution of IML PV neurons, coupling characteristics of AACs were not compared between control and post-SE mice. Together, these data demonstrate a post-SE decrease in frequency of both excitatory and inhibitory synaptic inputs to AACs.

### AAC output to granule cells is reduced after SE

To determine if AAC regulation of GCs is altered after SE we examined AAC driven synaptic inputs to GCs. Since our *RNAScope* and morphological studies identified that majority of the IML PV are AACs (Figure 1), we targeted IML PV neurons in PV-ChR2 reporter mice for discrete single-neuron optical stimulation and recorded AAC mediated responses in GCs. Given the relatively sparse distribution of IML PV neurons (2-3 cells/ section), DMD-based single neuron stimulation using a 50 µm ROI over the somata of isolated eYFP labeled IML neurons was used to activate individual AACs (*see methods*). Optically evoked AAC-mediated unitary IPSCs (uIPSCs) in GCs were recorded by selectively stimulating a single presynaptic AACs in the IML expressing eYFP-ChR2 using DMD coupled to blue LED (447nm) illuminated for 5ms single pulse for 10 sweeps for every 10s (Figure 4A, B). Regions with more than one eYFP-ChR2 positive somata within 50 µm of each other in the IML and GCL were excluded. In order to minimize direct activation of PV terminals on postsynaptic GC, only GCs with somata >150 µm lateral to the optically activate AAC somata were included in the analysis. Optical activation of AACs at 1-2 mW (in ROI) evoked uIPSCs in synaptically connected GCs (Figure 4A). In control experiments, blue light activation of eYFP-ChR2 expressing AACs consistently elicited a single action potential (not shown). Additionally, blue light activation of somata lacking eYFP or amber light (589 nm) activation of eYFP +ve somata consistently failed to evoke responses in recorded GCs (Figure 4A). In a subset of connected AAC to GC pairs, the presynaptic AAC was patched, held in current clamp, and activated by brief current pulses (0.8 nA) for 5ms to evoke single action potentials (Figure 4C) to confirm both the presence of uIPSC in the synaptic coupled GC and cellular/axonal morphology of the AAC (n= 3 pairs). These extensive validation studies confirmed that we could reliably activate single ChR2 expressing AACs and record uIPSCs in GCs.

Since uIPSC parameters in response to optical and current clamp activation of AACs were similar, we pooled uIPSCs evoked by optical stimulation (17/20 pairs in control; 6/10 pairs post-SE) and current injections under whole cell configuration (3/20 pairs in control; 4/10 pairs in post-SE) for analysis. The average uIPSC peak amplitude in GCs in response to AAC activation, calculated by averaging 10 sweeps including failures, was significantly decreased after SE (Figure 4B-D, average uIPSC peak amplitude in pA, Control: 128.47±21.95, n=20; post-SE: 40.42±7.15, n=9; unpaired t-test p<0.05). Similarly, the uIPSC amplitude potency, calculated by averaging the successful postsynaptic events excluding failures was also decreased after SE (Figure 4E, uIPSC peak amplitude potency in pA, Control: 131.13±21.66, n=20; post-SE: 53.72±17.06, n=9; unpaired t-test p<0.05). However, the decay kinetics and success rate of uIPSCs at the AAC to GC synapse was not altered in post-SE mice (Figure 4F-G, uIPSC decay in ms: Control: 7.13±0.86; post-SE: 10.36±1.73; unpaired t-test p>0.05. P_success_: Control: 0.95±0.02, n=20; post-SE: 0.83±0.10, n=9; unpaired t-test p>0.05) indicating that the reduction was likely due to a decrease in postsynaptic receptors . The connection probability between AAC and GC showed a trend towards a decrease which did not reach significance (Figure 4H, probability of synaptic connections in % - Control: 50%; 25/50 pairs; post-SE: 32.3%; 10 /31 pairs; χ^2^ _(1, 81)_= 2.45, p = 0.12 chi-square test with Yates correction). IPSCs evoked by AAC activation at 20 Hz showed robust short depression in both control and post-SE mice (Figure 4I, J) which is consistent with the high release probability of at AAC synapses and is similar to what has been reported in PV-BC synapses (Hefft and Jonas, 2005; Yu *et al*., 2016a; Zhang and Buckmaster, 2009). Despite the decrease in peak amplitude of AAC evoked uIPSC in GC after SE (amplitude in pA, Figure 4I, Control _IPSC1_: 230.16±58.4, n=8 pairs; Post SE _IPSC1_: 77.08±27.2, n=6 pairs, between treatment, Two way ANOVA F(1,130) = 36.8, p<0.001), the rate of synaptic depression elicited by a 20Hz train was not different between groups (Figure 4K, ratio of amplitude of IPSC10 to IPSC1, Control_IPSC10/IPSC1_: 0.31 ±0.04, n=8 pairs; Post SE _IPSC10/IPSC1_: 0.27±0.04, n=6 pairs, between treatment p>0.05 by Two way RM ANOVA). Together, data demonstrate a significant weakening in potency of AAC to GC synapses and suggests that AAC mediated AIS inhibition is reduced early after SE.

### AIS GABA reversal is depolarizing and unchanged after SE

Previous studies have identified that the reversal potential for GABA currents (*E_GABA_*) at AAC synapses on AIS of cortical pyramidal neurons show a delayed developmental switch to hyperpolarization relative to the resting membrane potential (Rinetti-Vargas *et al*., 2017). However, whether *E_GABA_* in the AIS of GCs, which rest at relatively hyperpolarized membrane potentials, is depolarizing has not been examined. Moreover, whether *E_GABA_* at GC AIS is altered early after SE, as observed in the somato-dendritic compartments of GCs (Pathak *et al*., 2007), is currently unknown. We used gramicidin perforated-patch recordings to measure *E_GABA_* at GC AIS in brain slices from control and post-SE mice. Epifluorescence images were conducted during perforated patch recordings including both 5-CFDA, a dye permeable through the perforation, as well as Alexa 594 which is excluded from the perforated pore. As illustrated in Figure 5A, 5-CFDA (green) but not Alexa 594 (red) fluorescence was visualized in the somata upon establishment of the perforated patch. The appearance of Alexa 594 fluorescence marked the transition to whole cell configuration and was accompanied by a change in *E_GABA_* to 0 mV reflecting the high chloride internal solution (Supplementary Fig. 7). Muscimol (100 μM), a GABA_A_R agonist, was applied on to the GC AIS using a picospritzer to elicit GABA currents. Figure 5B illustrates the relative location of the recording electrode and the muscimol puff (stim) electrode. Note that the puff electrode targets the 5-CFDA labeled AIS (Figure 5B3) and elicited inward currents at the holding potential (V_hold_) lower than -70 mV (Figure 5C). GABA currents in response to muscimol application were recorded at various holding potentials and used to determine GABA reversal potential as the voltage at 0 current (Figure 5C, Supplementary Fig. 7A). Recordings were discontinued when Alexa 594 was visualized in the soma or if there was a sudden drop in access resistance. Control experiments confirmed that muscimol and not mechanical pressure contributed to the inward currents and that muscimol-evoked currents were blocked by the GABA_A_R antagonist SR95531 (10 µM) (Supplementary Fig. 7B-C). In recordings from control and post-SE slices, GC AIS *E_GABA_* was significantly depolarized compared to the resting membrane potential (Figure 5C-D, Ctrl: RMP = -88.39 ± 1.48 mV, *E_GABA_* = -65.59 ± 1.25 mV, n=6 cells from 3 mice, p < 0.0001; Post-SE: RMP = -85.50 ± 1.20 mV, *E_GABA_* = -60.96 ± 2.77 mV, n=8 cells from 4 mice p< 0.0001 by paired t-test). However, there was no difference in either AIS *E_GABA_* or resting membrane potential between GCs from control and post-SE mice (Figure 5D, Mdn^Ctrl^ = -65.88 mV, Mdn^Pilo^ = -61.82 mV, U = 15, p = 0.2824,). These data indicate that *E_GABA_* AAC to GC synapses are not altered after SE.

**Figure 5:**
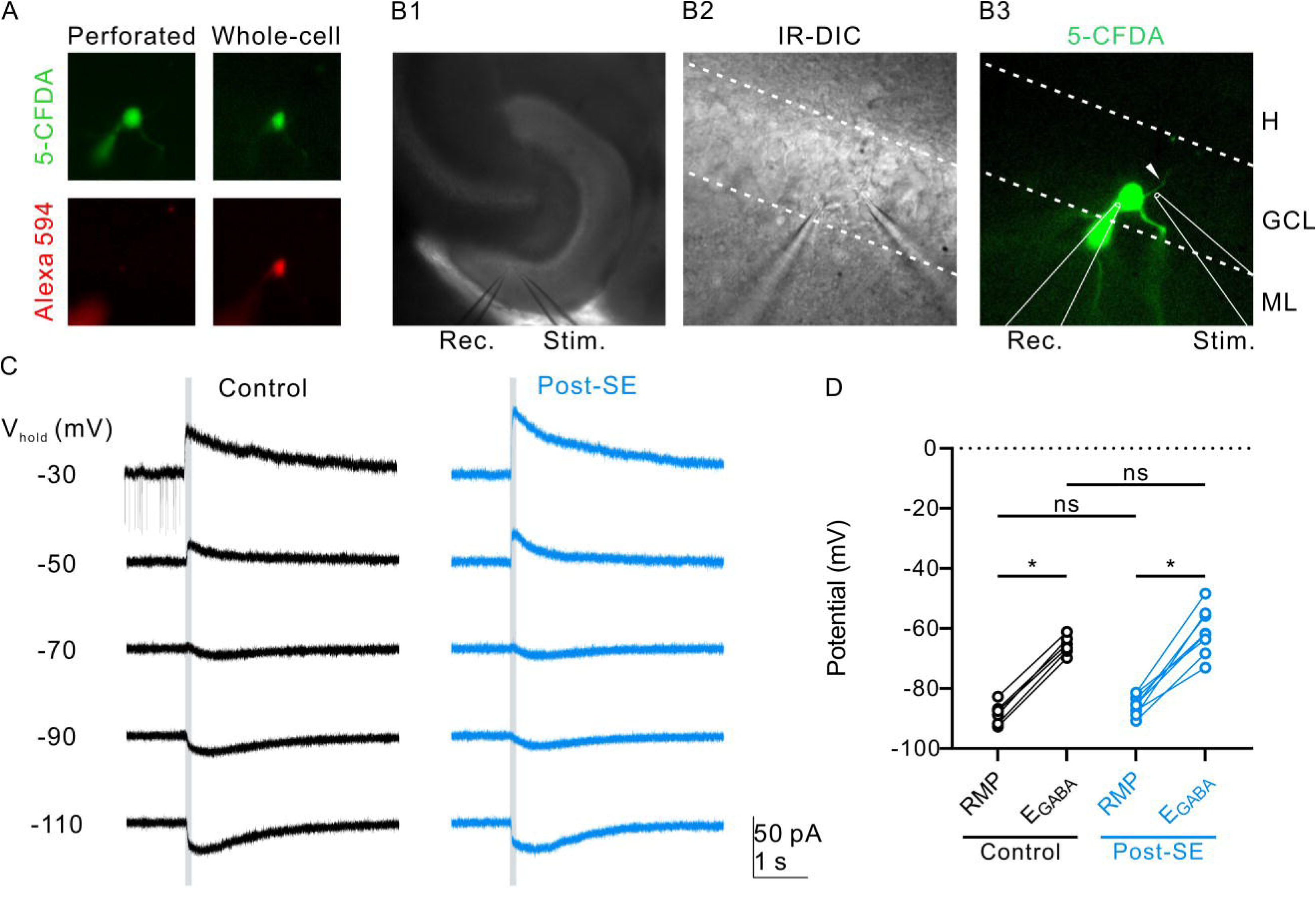
AIS-mediated GABA currents are depolarizing in both control and post-SE GCs. (A). Representative epifluorescence images obtained during gramicidin perforated-patch show confirmation of perforated patch mode by presence of 5-CFDA and absence of Alexa 594 signal in the recorded GC (on the left). Somatic Alexa 594 fluorescence on the right illustrates the rupture of perforated patch and entry into whole cell mode. (B) Low-magnification (B1), high- magnification IR-DIC (B2), and epifluorescence (B3) images illustrate placement of the stimulation pipette (Stim.) targeting the axon initial segment (AIS, arrowhead) and patch pipette (Rec.) at the GC somata. (C) Gramicidin-perforated patch recordings of muscimol-evoked GABA currents in the recorded GC during muscimol stimulation of AIS in control (left) and post-SE (right) mice. The holding potentials (V_hold_) are indicated on the left. (D) Summary of the resting membrane potential (RMP) and equilibrium potential of GABA (*E_GABA_*) in AIS in control and post-SE mice. *P < 0.0001, two-tail paired t-test.

### GABA_A_ receptors at AAC synapses on to GCs are reduced after SE

The selective reduction in amplitude potency at AAC synapses on GCs without reduction in release probability suggests a decrease in postsynaptic GABA_A_ receptors. Using peak-scaled nonstationary fluctuation analysis (Sigworth, 1980; Traynelis *et al*., 1993; Yu *et al*., 2016b), we evaluated the change in single channel current and receptor number at AAC to GC synapses. Although the peak amplitudes were much lower and less variable in synapses from post-SE mice, the characteristic parabolic distribution of mean-variance data was observed at AAC to GC synapses from both control and post-SE mice (Figure 6A). Single channel current estimates between synapses in control and post-SE mice were not different (Figure 6B, in pA, Control: 1.24±0.36, n=13; post-SE: 1.3±0.27, n=7, by Mann Whitney U-test p>0.05), indicating that GABA_A_ receptor conductance is unchanged following SE. However, the number of channels open at peak in the post-SE group was significantly reduced to approximately half that of controls (Figure 6C, Control: 110.97±16.8, n=13; post-SE: 51.56±14.6, n=7, by Mann Whitney U-test p<0.05). These findings indicate a selective weakening of AAC synapses on GCs early after SE.

**Figure 6:**
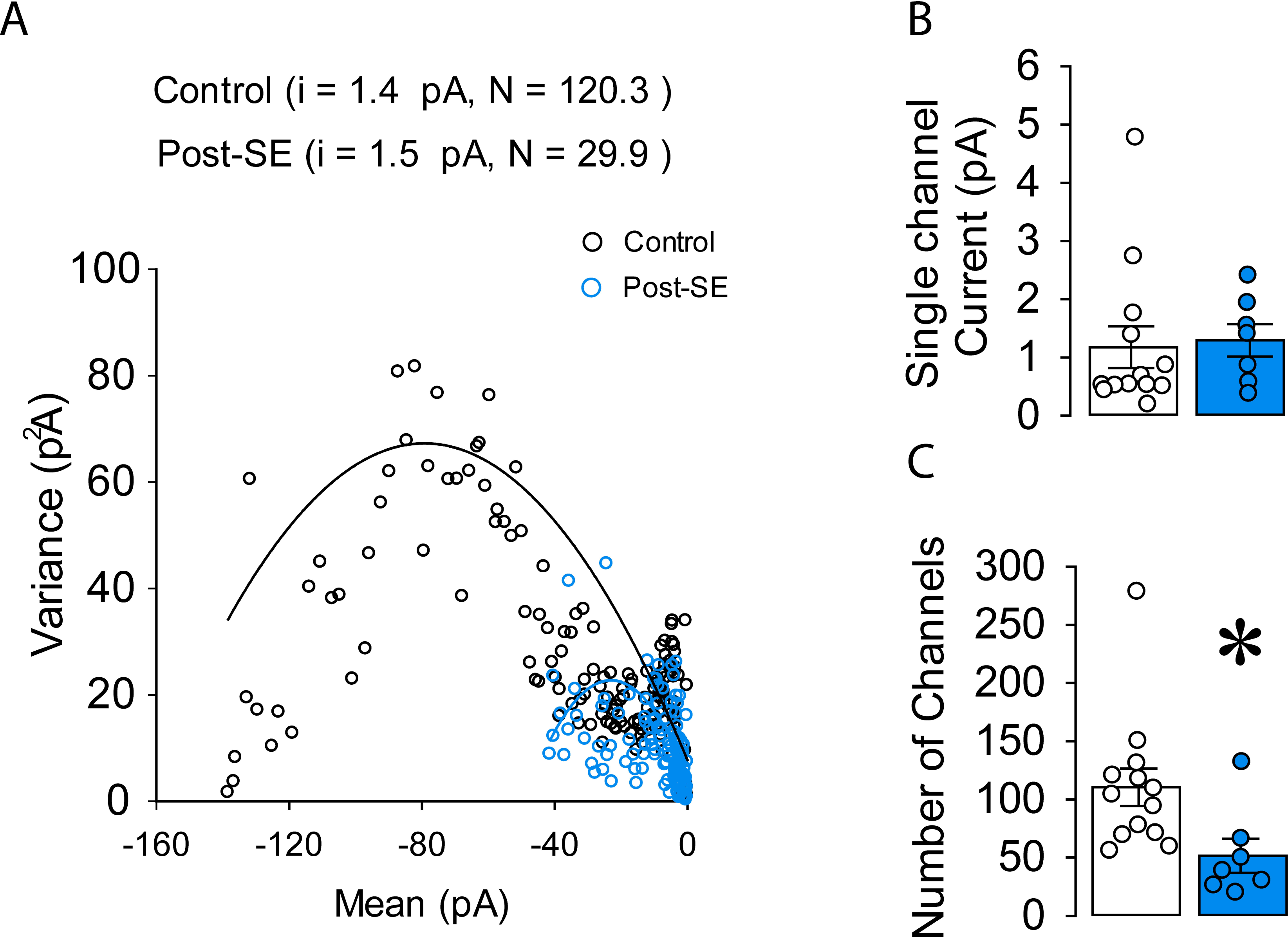
Decrease in postsynaptic GABA_A_ receptors at AAC-GC synapses. (A) Summary mean-variance current plots from control (black) and post-SE (blue) obtained from uIPSCs at AAC-GC synapses. (B-C) Summary plots represent weighted-mean single-channel currents (B) and the number of channels open at the peak current (C) obtained from curves fitted with equation f(x) = ix –x2/N, where ‘i’ is weighted-mean single-channel current, ‘N’ is the number of channels open at the peak.

Inhibitory synapses on AIS express GABA_A_ receptor α2 subunits (Fritschy *et al*., 1998). To test whether SE leads to a reduction in putative postsynaptic sites with GABA_A_ receptor α2 subunits at the AIS, we undertook immunostaining for GABA_A_ receptor α2 subunits and AIS using the AIS specific scaffolding protein Ankyrin G (Gao and Heldt, 2016; Jenkins and Bennett, 2001). While AIS length remained unchanged after SE (Supplementary Fig. 8A-C, in µm, Control: 40.75±4.22, n=8 segments/5 mice; post-SE: 42.39±4.65, n=12 segments/5 mice by unpaired t-test), the number of putative postsynaptic sites labeled for GABA_A_ α2 subunits located in close apposition to the AIS was decreased (Supplementary Fig. 8D, Control: 14.50±2.33, n=8 segments/5 mice; post-SE: 9.00±2.17, n=12 segments/5 mice by unpaired t-test, p<0.05). Correspondingly, the density of sites labeled for GABA_A_ receptor α2 subunits found in close apposition to the AIS was also reduced (Supplementary Fig. 8E, Control: 0.356±0.0590, n=8 segments/5 mice; post-SE: 0.212±0.0591, n=12 segments/5 mice by unpaired t-test, p<0.05).

### Post-SE reduction in synaptic strength compromises AAC dependent shunting inhibition of GC firing

The reduction in the number of putative AIS synapses labeled for GABA_A_ receptor α2 subunits density after SE, identified in our immunofluorescence, and reduction in receptor numbers in nonstationary fluctuation analysis, indicate compromised GABA conductance. Based on the experimental GABA reversal potential, the single channel conductance, and channel numbers estimated from nonstationary fluctuation analysis we calculated the peak GABA conductance at an AAC to GC synapse at the AIS to be approximately 4.8nS in controls and 1.48nS after SE. We performed simulations on a multicompartmental GC model (Santhakumar *et al*., 2005; Yu *et al*., 2016a) and included an AIS compartment (Figure 7A-B, also see methods) to examine the efficiency of GABA conductance in shunting synaptically driven GC action potential firing. Model GC was activated with increasing levels of peak excitatory AMPA conductance (low:14 nS – high:30 nS, Figure 7C) at the two simulated medial perforant path inputs to the granule cell. The effect of increasing GABA conductance at each of the two simulated AAC to AIS synapses from 0.5 to 10 nS (with decay set at 7 ms based on uIPSC decay in controls), on GC firing was examined (Figure 7D). GABA conductance in the range observed in controls (4.5-5.5 nS) was effective in shunting action potential firing for a range of AMPA currents and were permissive to firing when the AMPA conductance was increased. In contrast, GABA conductance in the range observed in post-SE AIS synapses failed to suppress firing even in response to low levels of excitation. These findings demonstrate that the post-SE change in conductance compromises AAC-mediated shunting of GC firing (Figure 7D, E). Although the decay kinetics of uIPSCs was not different between groups (Figure 4F), decay time in synapses from post-SE GCs trended to be higher (about 10 ms). Since synaptic decay kinetics can have a significant impact on GABAergic inhibition, we performed additional simulations using the AMPA and GABA conductance parameter space above with GABA synaptic conductance set to 10 ms to better simulate the post-SE condition. Even with the slower decay, the GABA synapses constrained by the conductance values recorded in GC from post-SE mice were less efficacious at suppressing afferent driven GC firing than synapses with 7 ms decay and constrained by the conductance values observed in controls (comparing Figures 7 D and F). These simulation studies indicate that the post-SE reduction in AAC synaptic strength could comprise overall AAC dependent inhibition at GC synapses.

**Figure 7:**
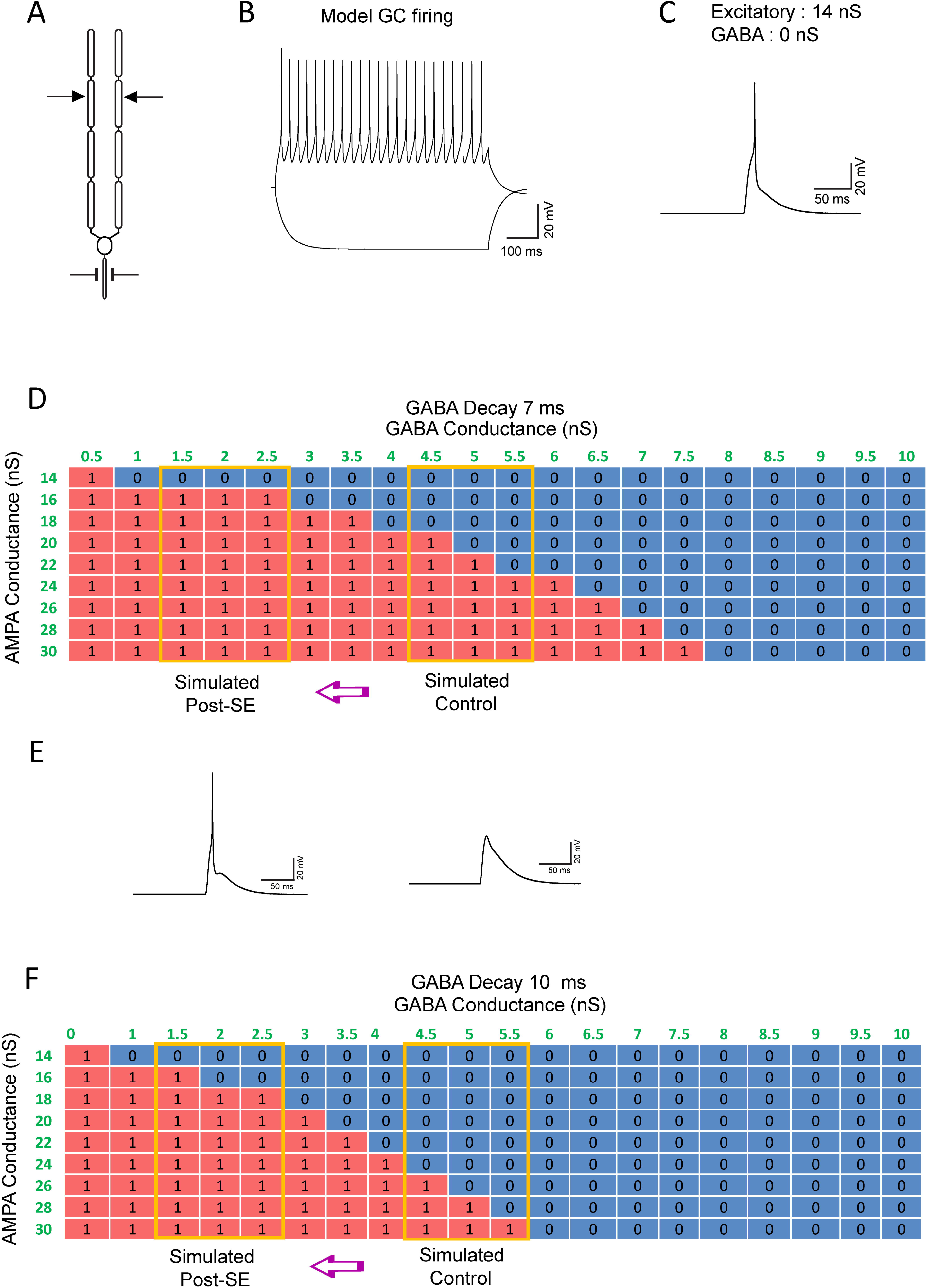
Post-SE changes in GABA conductance compromises AAC dependent shunting inhibition at AIS synapses. (A) Schematic of multicompartmental model GC showing location of excitatory synapses in the middle molecular layer compartment of the two dendrites and inhibitory synapse location on AIS. (B) Characteristic voltage responses of model GC during +300pA and -200pA current steps. (C) Model GC firing in response activation of excitatory conductance in the MML and in the absence of inhibitory conductance at the AIS segment. (D) Summary heatmap with GABA decay set to control mean value of 7 ms, shows GC firing (1) in response to increasing AMPA and GABA conductances. The range representing estimated GABA conductance in control and post-SE GCs is boxed in yellow. 1 – Represents action potential (red) and 0 – represents shunting inhibition of firing (blue). (E) Representative action potential (left) and EPSP (right) illustrate the simulated membrane traces with firing indicated as 1 in heat map and lack of action potential denoted by 0 in the heat maps. (F) Summary heatmap with GABA decay set to 10 ms, the value observed in AAC to GC synapses in post-SE mice. GC response to increasing AMPA and GABA conductances are as in D.

## Discussion

Combining colocalization of parvalbumin and PTHLH, a reliable marker for axo-axonic cells (Paul *et al*., 2017), and neuronal morphology, the current study identifies that PV neurons in the dentate IML are predominantly axo-axonic cells providing a viable approach to target this sparse neuronal population for physiological analysis. Despite sharing high-frequency, non-adapting firing characteristics and low input resistance with PV-BCs, AACs have a significantly higher rheobase for activation and a more depolarized action potential threshold. Although AACs need greater depolarization to reach threshold, their maximum firing rate trended to be higher than in PV-BCs. Unlike PV-BCs, AACs lacked persistent firing upon repolarization. Thus, the intrinsic physiology of dentate AACs is ideally suited to respond to strong sustained increases in network activity with high-fidelity firing. Our analysis revealed that the frequency of spontaneous excitatory and inhibitory synaptic currents was reduced after SE. Moreover, we identified a significant decrease in maximum firing frequency of AACs after SE which was associated with an increase in membrane capacitance. Simultaneously, although GABA reversal potential at AIS synapses, uIPSC reliability, and short-term plasticity of AACs to GCs synapses remained unchanged, the uIPSC amplitude was significantly decreased after SE. Non-stationary noise analysis was consistent with a decrease in GABA_A_ receptors at AIS synapses. Additionally, there was an apparent reduction in the number of postsynaptic sites immunolabeled for GABA_A_R α2 subunits at the GC AIS after SE. To our knowledge, our study is the first to evaluate functional changes in AAC in a model of experimental epilepsy and demonstrates a reduction in synaptic input, intrinsic firing, and synaptic output in dentate AACs after SE. Computational simulation of the effect of AIS synapses on synaptically evoked GC firing demonstrated a loss in the efficacy of shunting inhibition in models constrained by the post-SE AIS GABA conductance. Taken together, these changes indicate a diminished role of AACs in the dentate circuit early after status epilepticus. This functional uncoupling of AACs from the epileptogenic network and loss in inhibitory efficacy at the AIS could undermine an essential break to dentate throughput and contribute to epileptogenesis.

### Localization and synaptic physiology of dentate AACs

Since the early studies characterizing their extensive and unique axonal targeting synapses (Buhl *et al*., 1994; Somogyi *et al*., 1982; Szentagothai and Arbib, 1974), there has been great interest in understanding AACs regulation of neuronal output, brain rhythms and their role in disease (Dudok *et al*., 2022; Gallo *et al*., 2020; Howard *et al*., 2005). Recognition that labeling a cohort of inhibitory neuron precursors expressing the homeobox protein Nkx2.1 at a late embryonic stage selectively labels cortical AACs (Taniguchi *et al*., 2013) spearheaded the neurochemical and physiological analysis of cortical AACs (Gallo *et al*., 2020; Ishino *et al*., 2017; Paul *et al*., 2017). Although the labeling strategies used to identify cortical AACs fail to label hippocampal AACs, single-cell sequencing of cortical AACs has yielded specific markers that have been leveraged to target AACs in the hippocampal CA1 (Dudok *et al*., 2021; Ishino *et al*., 2017; Paul *et al*., 2017). Here we show, for the first time, that PTHLH also labels PV+ve AACs in the dentate IML. Consistent with earlier studies using Golgi staining (Soriano and Frotscher, 1989; Soriano *et al*., 1990), we show that the morphology of PV neurons in the dentate IML is consistent with AACs. The multipolar dendritic morphology with prominent apical dendrites and somatic location at the border of the GCL and molecular layer of AACs in our study is consistent with the structure and layered location pattern in cortical AACs (Taniguchi *et al*., 2013). Our finding also complements earlier studies which tracked the somata of cells extending axonal cartridges to GC AIS to PV+ve neurons in the dentate molecular layer in rat (Soriano *et al*., 1990). In keeping with the presence of morphologically identified AACs in the GCL and hilus (Elgueta *et al*., 2015; Overstreet and Westbrook, 2003), we report sparse PV and PTHLH labeled, presumed AACs in these regions. A subset of PTHLH-positive neurons failed to localize PV suggesting that, similar to the cortex (Paul *et al*., 2017), DG may have more than one neurochemical AAC subtype. These data indicate that IML PV-INs can be reliably targeted for functional analysis of dentate AACs.

Available physiological data from dentate AACs suggest that they share intrinsic membrane properties and fast-spiking characteristics with PV-BCs (Buhl *et al*., 1994; Overstreet and Westbrook, 2003). Our comparison identified that AACs need greater depolarization to reach threshold but show more reliable high-frequency firing during depolarization and lack the persistent discharges in the absence of depolarization observed in BCs (Elgueta *et al*., 2015). Indeed, despite sharing fast spiking characteristics, cortical BC and AACs have been found to differ in certain intrinsic properties (Taniguchi *et al*., 2013). Moreover, AACs also show regional differences in physiology, morphology, and expression of neurochemical markers (Buhl *et al*., 1994; Ishino *et al*., 2017). Apart from similarities in intrinsic physiology, AAC synapses on GCs have high success rates and robust paired pulse and short-term depression (Figure 4), as seen in BC to GC synapses (Bartos *et al*., 2002; Overstreet and Westbrook, 2003; Yu *et al*., 2016a). Together these physiological features of dentate AACs and their extensive apical dendritic arbors, make them ideally suited to sense afferent excitability, respond reliably during high input activation, and inhibit action potential initiation by projection neurons, as has been reported in response to whisker stimulation in the barrel cortex (Zhu *et al*., 2004). Thus, AACs are in a privileged position to support the dentate inhibitory gate and limit dentate throughput during re-entrant seizure-like activity.

### Seizure-induced changes in AAC intrinsic and synaptic physiology

Given the potential for AACs to clamp down action potential initiation (Schmidt-Hieber and Bischofberger, 2010) and the role for altered dentate inhibition during epileptogenesis, several studies have examined histopathological changes in dentate AAC to GC synapses. However, functional analysis of AACs in a model of epilepsy has been lacking (Dudok *et al*., 2022). Studies in human epileptic tissue have found that although PV-INs in the hilus are reduced, the PV-IN density in the dentate IML remains unchanged (Wittner *et al*., 2001). Despite the caveat that PV immunoreactivity can be altered in disease, our data showing maintenance of PV-INs in IML after pilocarpine-SE are consistent with the human epilepsy data (Wittner *et al*., 2001) and suggest that AACs largely survive SE induction. Several studies have reported loss of AAC terminals in human epileptic circuits with the most profound loss observed in the DG (Arellano *et al*., 2004; Ribak, 1985), while others have reported an increase in PV synapse on the AIS (Wittner *et al*., 2001). A recent study using a combination of markers to identify AAC synapses found an increase in PV-BC synapses and a decrease in AAC synapses in the human epileptic dentate gyrus (Alhourani *et al*., 2020; Lillis, 2020), suggesting a selective reduction in AAC mediated inhibition in epilepsy. Our direct assessment of AAC physiology after experimental SE identified that, despite a lack of change in passive membrane properties, there is a striking reduction in the high frequency firing of AACs after SE. Further analysis identified a reduction in the integrity of perineuronal nets and a corresponding increase in AAC capacitance after SE which could contribute to the decrease in AAC firing. Indeed, previous studies have shown that seizure-induced degradation of the perineuronal net can increase PV neuron capacitance and reduce intrinsic excitability without changes in action potential threshold (Tewari *et al*., 2018). Since our recordings were conducted in the absence of synaptic blockers, it is possible that changes in feedback synaptic inhibition or tonic GABA currents could also contribute to the post-SE decrease in AAC firing. These findings in AACs are distinct from the lack of change in the intrinsic physiology of PV-BCs after SE (Yu *et al*., 2016a) and would directly undermine activity-dependent GABA release at the synapses. The post-SE reduction in the frequency of sEPSCs and sIPSCs that we report in AACs parallels what has been reported in PV-BCs (Yu *et al*., 2013; Yu *et al*., 2016a; Zhang and Buckmaster, 2009). The reduction in basal synaptic input to AACs likely reflects the loss of hilar mossy cells and interneurons after SE (Buckmaster *et al*., 2017; Buckmaster and Jongen-Relo, 1999; Obenaus *et al*., 1993; Zhang *et al*., 2009). Additionally, it is possible afferent inputs to AACs are also reduced which could limit the recruitment of AAC mediated feedforward inhibition. While sIPSC amplitude showed a slight reduction, sEPSC amplitude was increased after SE suggesting potential compensatory regulation, as the average EPSC charge transfer over time was not different between control and post-SE groups (Supplementary Figure 5).

Our identification that AAC synapse on to GC show a profound decrease in unitary IPSC amplitude without a change in success rate after SE, differs from the increase in failures and amplitude of PV-BC synapses on GCs early after SE (Zhang and Buckmaster, 2009). Additionally, there was a trend towards a decrease in synaptic connectivity between AACs and GCs after SE which points to a further reduction in the ability of AACs to influence GC output. Thus, while the input received by PV-BCs and AACs is similarly decreased early after SE, AACs show a unique reduction in intrinsic firing and strength of synaptic output to GCs. Further evaluation of the GABA_A_ receptor expression using nonstationary noise analysis identified that a decrease in GABA_A_ receptors contributed to post-SE reduction in AAC synaptic strength. These changes, taken together, suggest functional uncoupling of AACs from the circuit which is likely to undermine their role in responding to heightened activity with robust synaptic inhibition of GCs.

### Plasticity of AIS Synapses

An intriguing aspect of AACs mediated inhibition is the possibility that AAC synapses can be depolarizing long after somatic and dendritic inhibition undergoes the developmental switch to hyperpolarization (Pan-Vazquez *et al*., 2020; Rinetti-Vargas *et al*., 2017; Szabadics *et al*., 2006). While it has been speculated that GABA may be depolarizing at the AIS (Szabadics *et al*., 2006; Woodruff *et al*., 2009), GABA reversal in cortical neurons has been shown to be hyperpolarizing at the age-range (>6 week postnatal) used in or study (Rinetti-Vargas *et al*., 2017). We find that while AIS GABA reversal (-60 mV) is more depolarized than the GC resting membrane potential (-80 mV), it is more negative than the action potentials threshold. Thus, AAC synapses would be expected to mediate shunting inhibition to GC (Schmidt-Hieber and Bischofberger, 2010). Indeed, we find that AIS synapses constrained by experimental conductance and depolarizing GABA reversal effectively shunt GC firing for a range of excitatory input drive. However, unlike GCs and PV-BCs which show a depolarizing shift in E_GABA_ one week after pilocarpine SE (Pathak *et al*., 2007; Yu *et al*., 2013), both AIS GABA reversal and GC resting membrane potential remained unchanged after SE. The post-SE decrease in AAC synaptic strength may reflect a homeostatic compensation to limit the AIS depolarization. This possibility is supported by recent studies demonstrating that chemogenetic activation of AACs reduces IPSC amplitude at AIS synapses in immature cortex, when GABA is depolarizing, while it increases IPSC amplitude at AIS synapses in the mature cortex when GABA is hyperpolarizing (Pan-Vazquez *et al*., 2020). Structural changes in AIS length which can impact neuronal spike threshold (Harty *et al*., 2013; Liu *et al*., 2017; Wefelmeyer *et al*., 2015) were not observed in post-SE GCs. Immunostaining for GABA_A_ receptor α2 subunits revealed potential reduction in colocalization with the AIS indicating a potential decrease in the number of AIS synapses. Since confocal imaging adopted here lacks the resolution to quantify receptor numbers, future studies using electron microscopy or super-resolution imaging could determine whether the number of receptors at AIS GABA synapses are reduced after SE. These findings together with the results of non-stationary fluctuation analysis indicate weakening of the AAC to AIS synapse after SE.

### Implications for epilepsy

Our findings demonstrate an early reduction in AAC excitability and its ability to inhibit GCs after SE. This reduction in inhibition at the site of action potential initiation is expected to compromise the strong inhibitory gating of the dentate gyrus which is critical for limiting dentate throughput and the spread of seizures (Carlson and Coulter, 2008; Dengler and Coulter, 2016; Krook-Magnuson *et al*., 2015; Krook-Magnuson *et al*., 2013). Additionally, activity sparsening mediated by dentate inhibition is also critical for pattern separation, a dentate dependent ability to discriminate patterns of neuronal activity representing overlapping behaviorally relevant information (Cayco-Gajic and Silver, 2019) which is compromised in epilepsy (Kahn *et al*., 2019; Madar *et al*., 2021). Moreover, hippocampal AACs have been found to contribute to maintaining location specific place cell firing indicating a role in memory processing (Dudok *et al*., 2021). Thus, the loss of AAC inhibition early after SE could contribute to both enhanced excitability and cognitive deficits associated with epilepsy. Interestingly, the advent of opto and chemogenetic approaches to manipulate specific neuronal populations has opened avenues for in vivo optical activation of parvalbumin neurons for seizure control (Krook-Magnuson *et al*., 2013; Paz *et al*., 2013). Since PV neurons include AIS targeting AACs, whether activating AACs which show differential plasticity in experimental epilepsy could paradoxically promote excitability needs to be considered. In this regard, the variable outcomes with optogenetic activation of PV neurons suppressing seizures in some studies (Ellender *et al*., 2014; Krook-Magnuson *et al*., 2013) while exacerbating it in others may be explained by opposite circuit effects of BCs and AACs after SE. Selective opto/chemogenetic manipulation of dentate AACs, needed to directly evaluate the role of AACs epileptogenesis, are currently limited by the low specificity of approaches to label dentate AACs (Ishino *et al*., 2017). Here we overcome these limitations by using a combination of PV labeling and layer specific localization to target AACs for functional analysis in a model of experimental epilepsy. Our study provides the first systematic physiological assay of dentate AACs and demonstrates a reduction in AAC excitability, synapse potency and AIS inhibition after SE. This undermining of AIS inhibition early after SE could augment dentate excitability and contribute to epileptogenesis.

## Supporting information

Proddutur-Supplementary Materials

## Acknowledgements

We thank Dr. David Carter at the UCR Microscopy Core for help with imaging. We thank Ds. Khaleel A. Razak and Jamiela Kokash for help with reagents and protocols for PNN staining.

## Author Contributions

A.P., C-W. Y., S.N., and A.G. performed experiments; A.P., S.N., C-W. Y., and A.G. analyzed data; A.P., S.N., C-W. Y., and V.S. interpreted results of experiments; A.P. C-W. Y., and S.N. prepared figures; A.P and V.S. conceived of and designed research V.S funding acquisition , A.P., S.N., C-W. Y., and V.S drafted manuscript.

## ABBREVIATIONS

AAC: Axo-Axonic Cell
aCSF: artificial Cerebrospinal Fluid
AHP: After Hyperpolarization
AIS: Axon Initial Segment
AP: Action Potential
BC: Basket Cell
ChR2: Channelrhodopsin2
DG: Dentate Gyrus
E_GABA_: GABA reversal potential
eYFP: enhanced Yellow Fluorescent Protein
GC: Granule Cell
GCL: Granule Cell Layer
IML: Inner Molecular layer
IQR: Inter Quartile Range
PNN: Perineuronal Net
PTHLH: Parathyroid Hormone Like Hormone
PV: Parvalbumin
PV-IN: Parvalbumin Interneuron
PV-BC: Parvalbumin Basket Cell
RMP: Resting Membrane Potential
R_in_: Input Resistance
SE: Status Epilepticus
sEPSC: spontaneous Excitatory Postsynaptic Current
sIPSC: spontaneous Inhibitory Postsynaptic Current
TLE: Temporal Lobe Epilepsy
uIPSC: unitary Inhibitory Postsynaptic Current
WFA: Wisteria Floribunda Agglutinin

## Supplementary Figures

**Supplementary Figure 1. Validation of transgenic mouse line.** Representative confocal images of PV immunolabeling on the left, eYFP expression the middle, and colocalization of PV and eYFP in the right. Dotted lines represent the contours of the dentate granule cell layer (GCL). IML: Inner Molecular layer. Scale bar: 100μm.

**Supplementary Figure 2. Example axonal cartridges from recorded AACs.** Representative confocal images of four (A-D) biocytin-filled axons recovered from PV neurons recorded in the IML shows distinct axonal cartridges (►) in the granule cell layer (GCL). The GCL is denoted by dotted lines. Single plane confocal images and have been rendered in black and white with an inverted color scheme for better visualization of axon cartridges. D_i_ and D_ii_ represent images obtained at 2 planes from the same cell. Scale bar: 50μm.

**Supplementary Figure 3. Quantification of NeuN and PV neurons following experimental status epilepticus in mice.** (A) Estimates of NeuN labeled cells in the hilus in the control and post-SE mice examined one week after SE induction. (B-C) Estimates of PV labeled neurons in Hilus (B) and inner molecular layer (IML in C) in control and post-SE mice. * Represents p<0.05 using unpaired t-test.

**Supplementary Figure 4. Quantification of WFA intensity peaks around IML PV neurons.**

(A) Confocal images of the immunofluorescence of PV (magenta) and WFA-labeled PNN (cyan) in PV neurons in the dentate inner molecular layer (presumed AACs) in control (A1) and post-SE (A2) mice. (B) Immunofluorescence intensity of WFA-labeled PNN in dentate AACs of control (B1) and post-SE (B2) animals. Color scale to the right shows increasing WFA intensity from black (0) to red (50%) with regions >50% intensity in white (>50-100%). WFA intensity peaks were quantified as the number of regions crossing the 50% intensity. (C) Summary quantification of WFA peaks normalized to ROI area. *p < 0.01, two-tail unpaired t-test. n=32 cells in Control group; n=33 cells in post-SE group from 5 mice each. Data are expressed as mean ± SEM. (D) Summary quantification of WFA peak count normalized to ROI area and averaged within animal. *p < 0.001, two-tail unpaired t-test. n=5 mice each. Data are expressed as mean ± SEM.

**Supplementary Figure 5. sEPSC and sIPSC kinetics In Control and Post-SE AACs** (A-D) Summary quantification of sEPSC weighted tau decay (A), rise time (B), charge transfer (C), and estimated charge transfer in one second (D) in control and post-SE cells. Estimated charge transfer in one second (D) was calculated as the product of the average EPSC charge transfer and frequency (E-H) Summary quantification of sIPSC weighted tau decay (E), rise time (F), charge transfer (G), and estimated charge transfer over one second (H) in control and post-SE AACs.

**Supplementary Figure 6. Gap junctional coupling in AAC-AAC neurons.**

Representative traces of subthreshold (top) and suprathreshold (bottom) membrane voltage from dual AAC recordings show the presence of gap junction coupling between AACs in slices from control (left) and post-SE (right) mice. Arrowheads point to membrane depolarization in electrically coupled cell in the absence of direct current injection.

**Supplementary Figure 7. Validation of perforated patch recordings in GC** (A) Representative current trace (A1) and current-voltage relation (A2) of muscimol puff-evoked GABA currents at different holding potentials in recordings conducted in gramicidin perforated-patch (filled circle) and whole-cell (open circle) configuration. (B) Example Muscimol- and aCSF-evoked responses in the same recorded neuron illustrate a lack of mechanical artifact. (C) Example trace (C1) and plot of normalized response vs. time (C2) illustrate the effect of GABA_A_R antagonist SR95531 (100 µM) on muscimol puff-evoked GABA currents. Note stable access resistance (Ra) despite decrease in muscimol evoked current in the presence of SR95531.

**Supplementary Figure 8. Reduction in GABA_A_A** _α_**2 receptor immunolabeling at AIS segments in post-SE mice**. (A-B) representative confocal images of sections immunolabeled for ankyrin-G to localize AIS (green) with GAB_A_A α2 receptor (red) in control (A) and Post-SE (B) and zoomed in images are shown in the middle panels respectively. The far-right panels show redrawn AIS segments in cyan and the colocalized puncta tagged with white circular markers. Note that since the puncta were visualized in 3D stacks some puncta cannot be seen in the 2D images. (C-E) Summary plot of AIS length (C), number of GAB_A_A α2 subunits (D) and density of GAB_A_A α2 subunits (E).

