## Supplementary material for "RECLUSIVE CHANDELIERS: FUNCTIONAL ISOLATION OF DENTATE AXO-AXONIC CELLS AFTER EXPERIMENTAL STATUS EPILEPTICUS": Proddutur-Supplementary Materials

### Proddutur et al Supplementary Materials

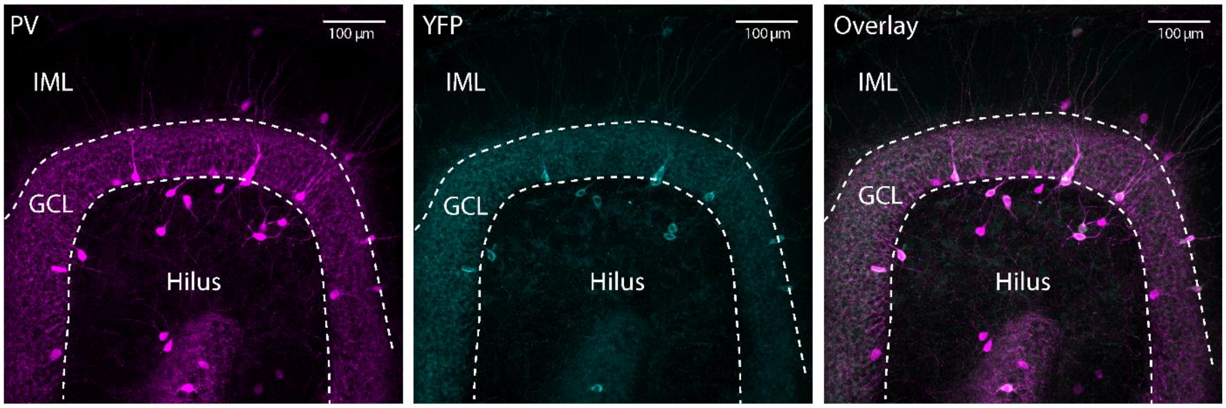


**Supplementary Figure 1. Validation of transgenic mouse line.** Representative confocal images of PV immunolabeling on the left, eYFP expression the middle, and colocalization of PV and eYFP in the right. Dotted lines represent the contours of the dentate granule cell layer (GCL). IML: Inner Molecular layer. Scale bar: 100µm.

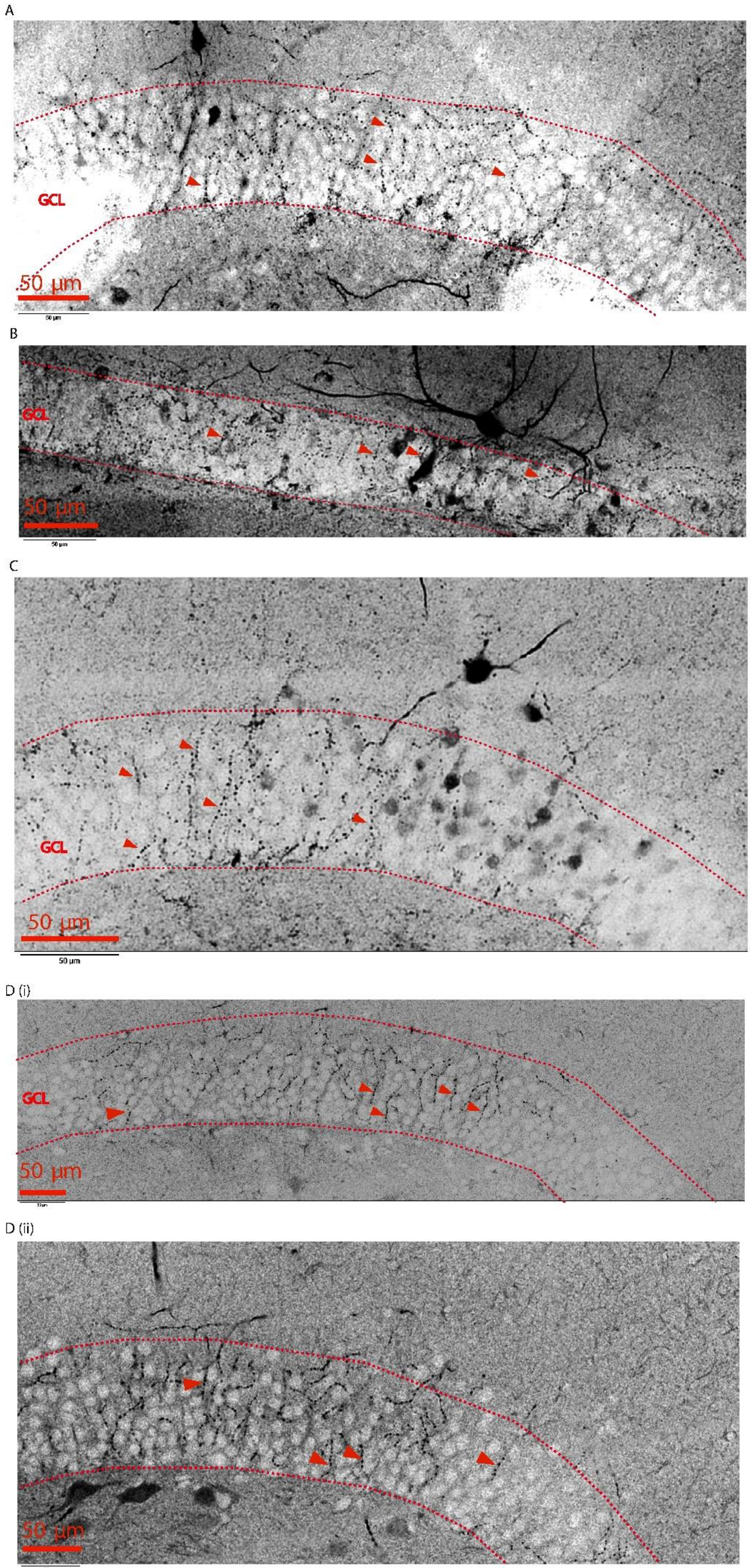

A
GCL
50 µm
B
GCL
50 µm
C
GCL
50 µm
D (i)
GCL
50 µm
D (ii)
50 µm

**Supplementary Figure 2. Example axonal cartridges from recorded AACs.** Representative confocal images of four (A-D) biocytin filled axons recovered from PV neurons recorded in the IML shows distinct axonal cartridges (►) in the granule cell layer (GCL). The GCL is denoted by dotted lines. Single plane confocal images and have been rendered in black and white with inverted color scheme for better visualization of axon cartridges. $D_i$ and $D_{ii}$ represent images obtained at 2 planes from the same cell. Scale bar: 50µm.

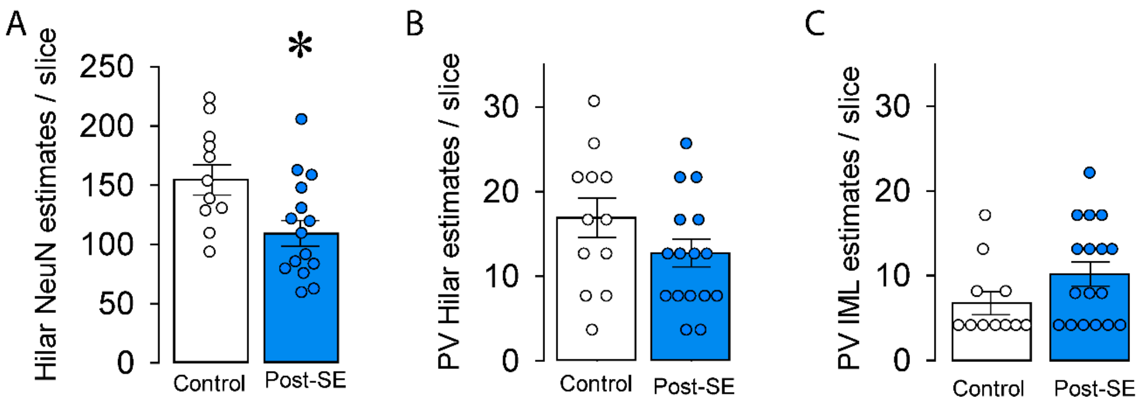


**Supplementary Figure 3. Quantification of NeuN and PV neurons following experimental status epilepticus in mice.** (A) Estimates of NeuN labeled cells in the hilus in the control and post-SE mice examined one week after SE induction. (B-C) Estimates of PV labeled neurons in Hilus (B) and inner molecular layer (IML in C) in control and post-SE mice. * Represents $p<0.05$ using unpaired t-test.

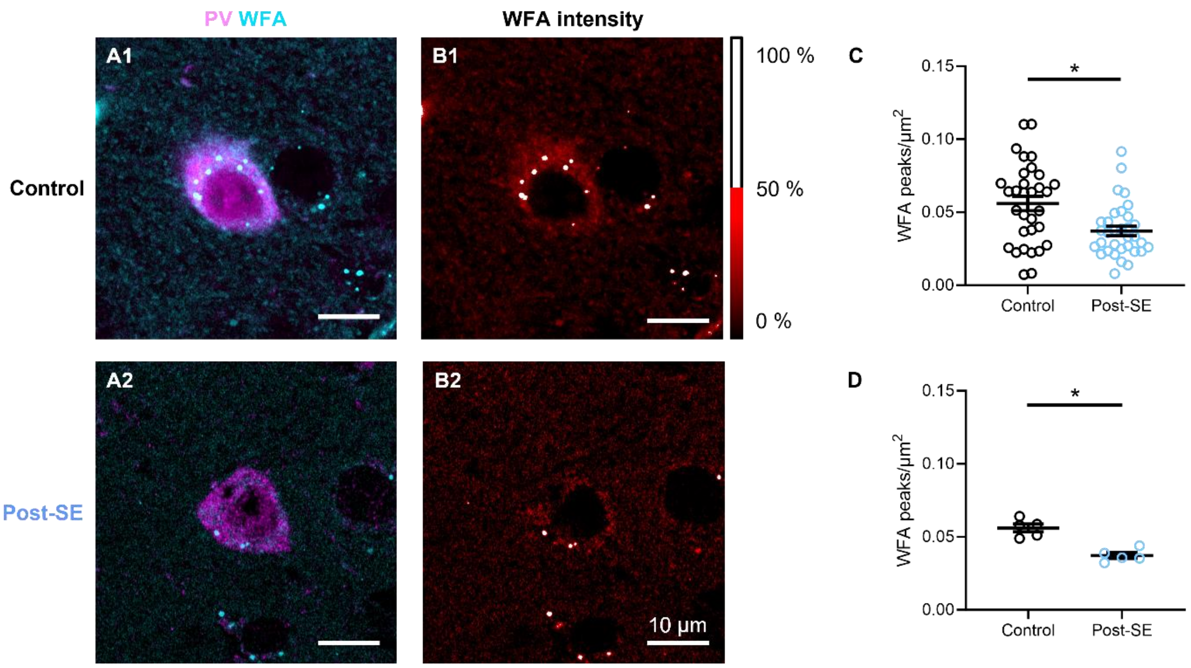


**Supplementary Figure 4. Quantification of WFA intensity peaks around IML PV neurons.** (A) Confocal images of the immunofluorescence of PV (magenta) and WFA-labeled PNN (cyan) in PV neurons in the dentate inner molecular layer (presumed AACs) in control (A1) and post-SE (A2) mice. (B) Immunofluorescence intensity of WFA-labeled PNN in dentate AACs of control (B1) and post-SE (B2) animals. Color scale to the right shows increasing WFA intensity from black (0) to red (50%) with regions >50% intensity in white (>50-100%). WFA intensity peaks were quantified as the number of regions crossing the 50% intensity. (C) Summary quantification of WFA peaks normalized to ROI area. *$p < 0.01$, two-tail unpaired t-test. n=32 cells in Control group; n=33 cells in post-SE group from 5 mice each. Data are expressed as mean ± SEM. (D) Summary quantification of WFA peak count normalized to ROI area and averaged within animal. *$p < 0.001$, two-tail unpaired t-test. n=5 mice each. Data are expressed as mean ± SEM

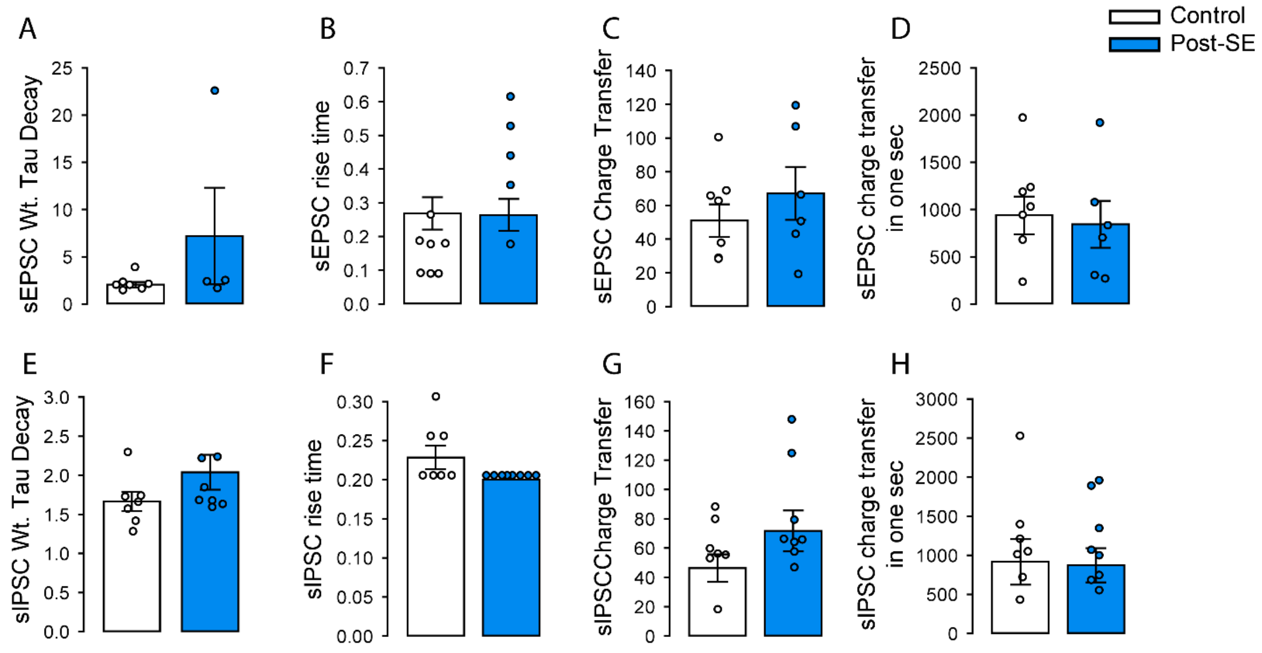


**Supplementary Figure 5. sEPSC and sIPSC kinetics In Control and Post-SE AACs** (A-D) Summary quantification of sEPSC weighted tau decay (A), rise time (B), charge transfer (C), and estimated charge transfer in one second (D) in control and post-SE cells. Estimated charge transfer in one second (D) was calculated as the product of the average EPSC charge transfer and frequency (E-H) Summary quantification of sIPSC weighted tau decay (E), rise time (F), charge transfer (G), and estimated charge transfer over one second (H) in control and post-SE AACs.

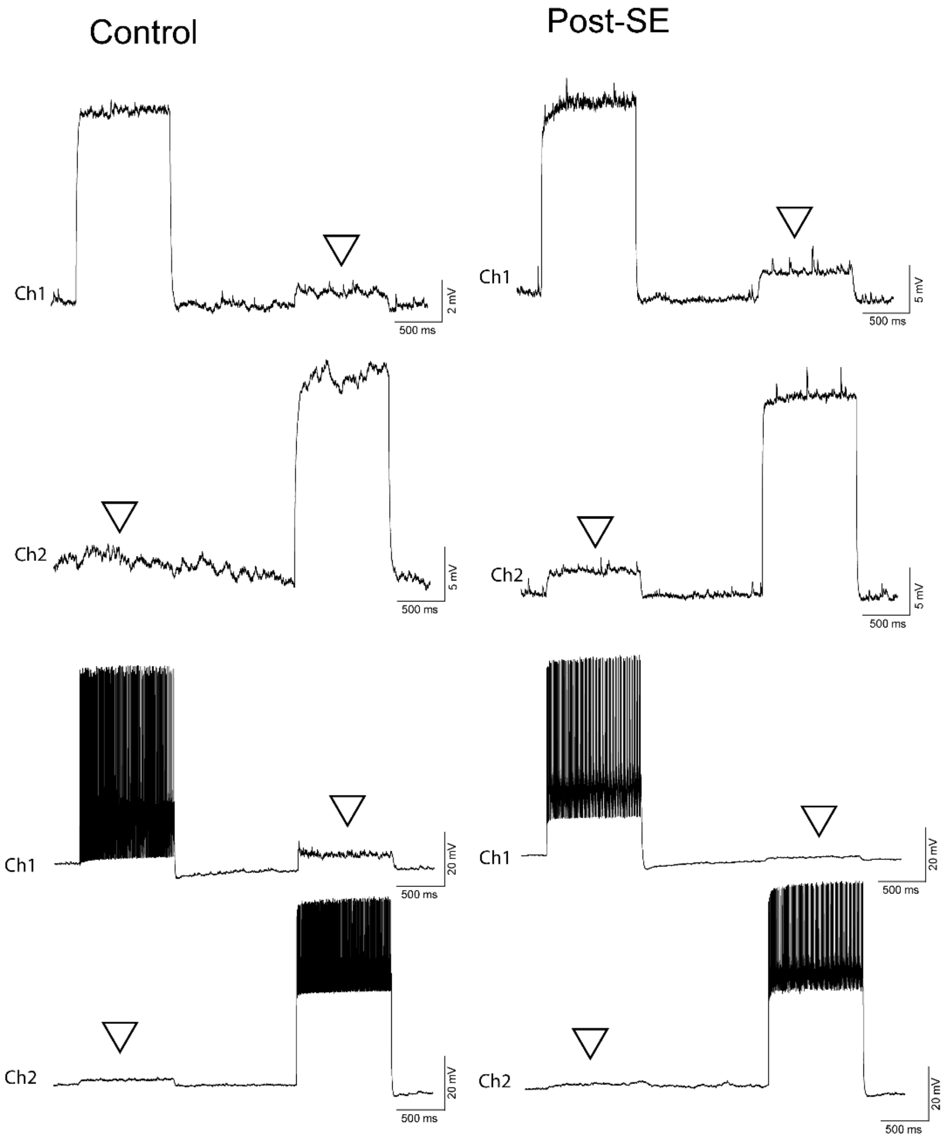


**Supplementary Figure 6. Gap junctional coupling in AAC-AAC neurons.**

Representative traces of subthreshold (top) and suprathreshold (bottom) membrane voltage from dual AAC recordings show the presence of gap junction coupling between AACs in slices from control (left) and post-SE (right) mice. Arrowheads point to membrane depolarization in electrically coupled cell in the absence of direct current injection.

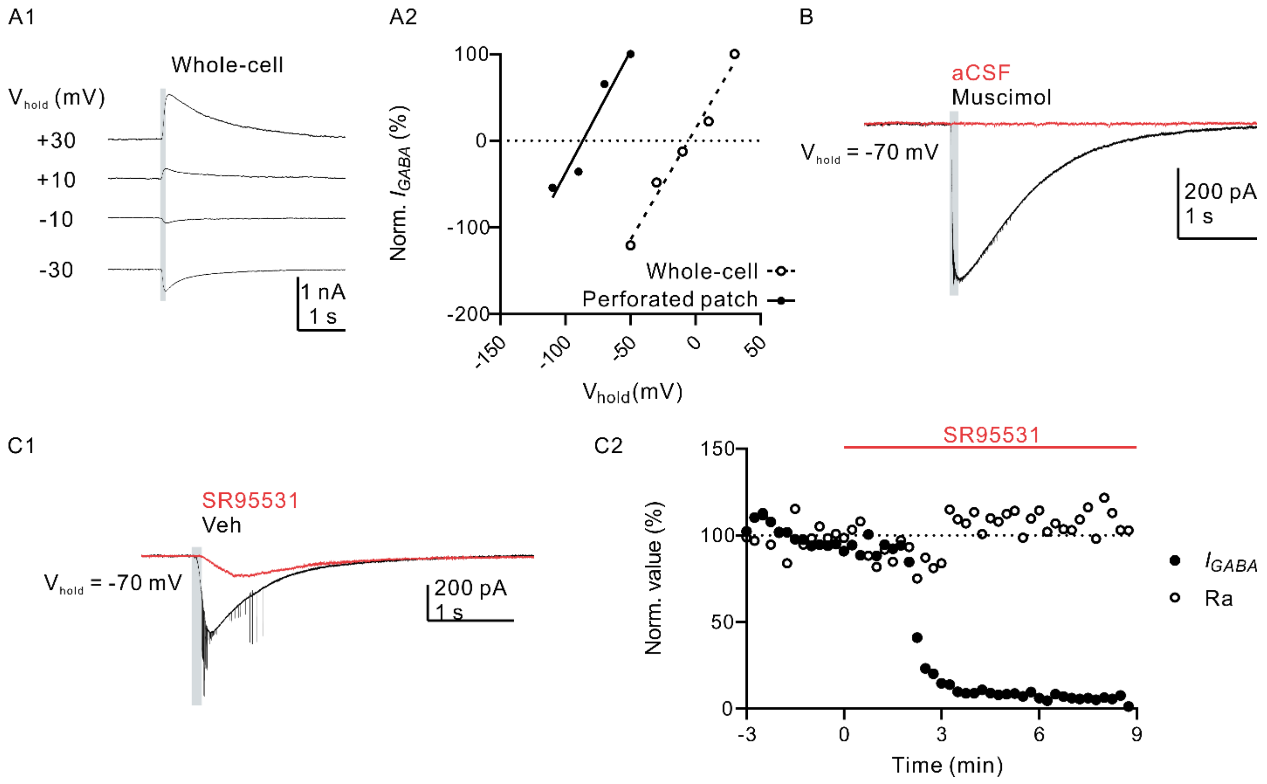


**Supplementary Figure 7. Validation of perforated patch recordings in GC** (A) Representative current trace (A1) and current-voltage relation (A2) of muscimol puff-evoked GABA currents at different holding potentials in recordings conducted in gramicidin perforated-patch (filled circle) and whole-cell (open circle) configuration. (B) Example Muscimol- and aCSF-evoked responses in the same recorded neuron illustrates lack of mechanical artifact. (C) Example trace (C1) and plot of normalized response vs. time (C2) illustrate the effect of $GABA_AR$ antagonist SR95531 (100 μM) on muscimol puff-evoked GABA currents. Note stable access resistance (Ra) despite decrease in muscimol evoked current in the presence of SR95531.

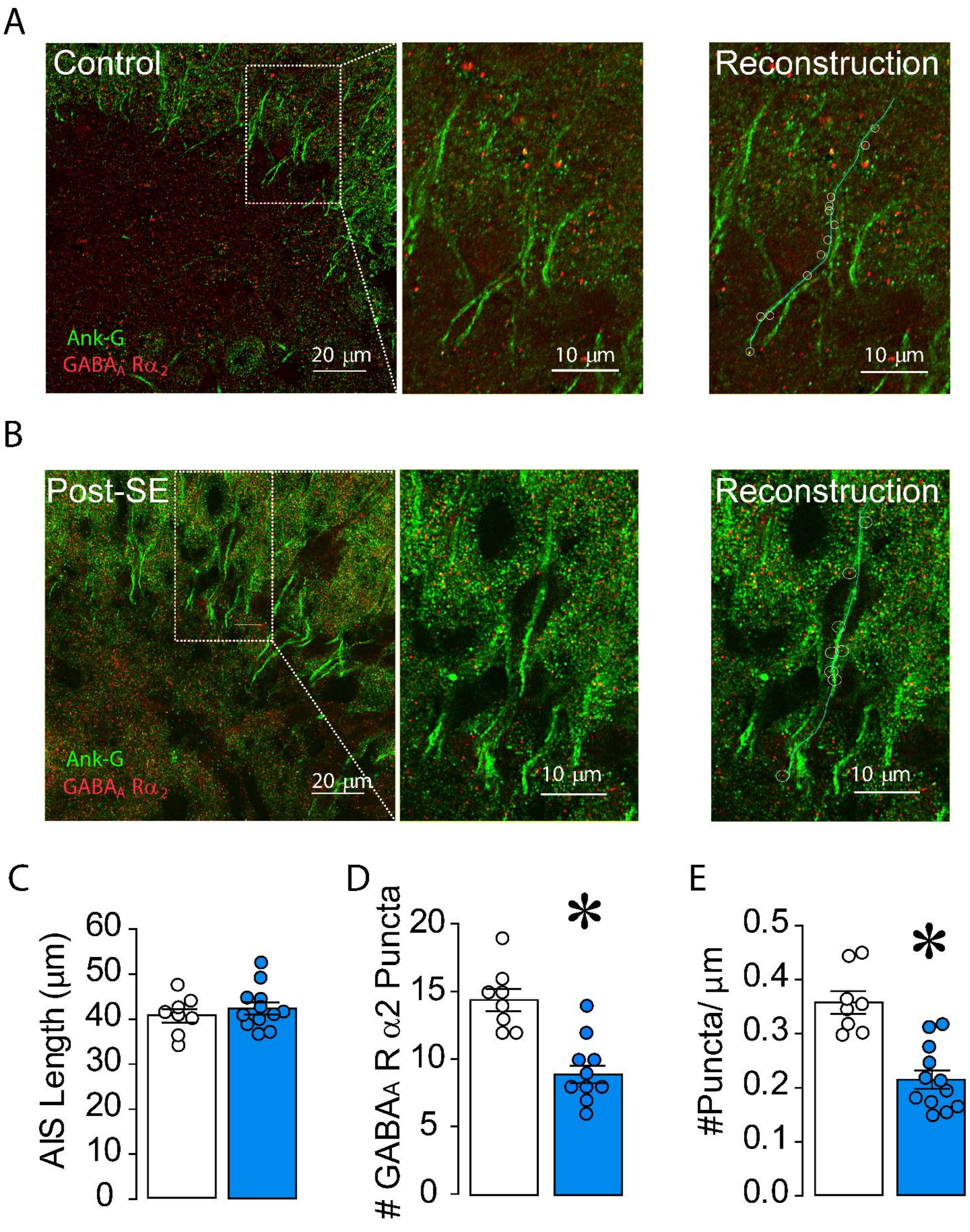
A
Control
Reconstruction
Ank-G
GABA$_A$ R$\alpha_2$
20 µm
10 µm
10 µm
B
Post-SE
Reconstruction
Ank-G
GABA$_A$ R$\alpha_2$
20 µm
10 µm
10 µm
C
AIS Length (µm)
0
10
20
30
40
50
60
D
### GABA$_A$ R $\alpha$2 Puncta
0
5
10
15
20
*
E
#Puncta/ µm
0.0
0.1
0.2
0.3
0.4
0.5
*

**Supplementary Figure 8. Reduction in GABA$_A$A α2 receptor immunolabeling at AIS segments in post-SE mice**. (A-B) representative confocal images of sections immunolabeled for ankyrin-G to localize AIS (green) with GAB$_A$A α2 receptor (red) in control (A) and Post-SE (B) and zoomed in images are shown in the middle panels respectively. The far right panels show redrawn AIS segments in cyan and the colocalized puncta tagged with white circular markers. Note that since the puncta were visualized in 3D stacks some puncta cannot be seen in the 2D images. (C-E) Summary plot of AIS length (C), number of GAB$_A$A α2 subunits (D) and density of GAB$_A$A α2 subunits (E).

| Parameter | Conductance (nS) | | | | | |
|---|---|---|---|---|---|---|
| | Soma | AIS | Dendrite in cell layer | Proximal Dendrite | Middle Dendrite | Distal Dendrite |
| Fast Na | 0.020 | 0.12 | 0.018 | 0.013 | 0.008 | 0.0 |
| Fast K | 0.016 | 0.018 | 0.004 | 0.004 | 0.001 | 0.001 |
| Slow K | 0.006 | 0.006 | 0.006 | 0.006 | 0.006 | 0.008 |
| Leak | 0.00004 | 0.00004 | 0.00004 | 0.000063 | 0.000063 | 0.000063 |

**Supplementary Table 1: Parameters modified from original model**
